# ATP hydrolysis tunes specificity of a AAA+ protease

**DOI:** 10.1101/2021.08.18.456811

**Authors:** Samar A. Mahmoud, Berent Aldikacti, Peter Chien

**Affiliations:** Department of Biochemistry and Molecular Biology, University of Massachusetts Amherst, Amherst, MA 01003, USA; Molecular and Cellular Biology Program, University of Massachusetts Amherst, Amherst, MA 01003, USA

## Abstract

In bacteria, AAA+ proteases such as Lon and ClpXP degrade substrates with exquisite specificity. These machines capture the energy of ATP hydrolysis to power unfolding and degradation of target substrates. Here, we show that a mutation in the ATP binding site of ClpX shifts protease specificity to promote degradation of normally Lon-restricted substrates. However, this ClpX mutant is worse at degrading ClpXP targets, suggesting an optimal balance in substrate preference for a given protease that is surprisingly easy to alter. *In vitro*, wildtype ClpXP also degrades Lon-restricted substrates more readily when ATP levels are reduced, similar to the shifted specificity of mutant ClpXP, which has altered ATP hydrolysis kinetics. Based on these results, we suggest that rates of ATP hydrolysis not only power substrate unfolding and degradation, but also tune protease specificity. We consider various models for this effect based on emerging structures of AAA+ machines showing conformationally distinct states.

Graphical Abstract

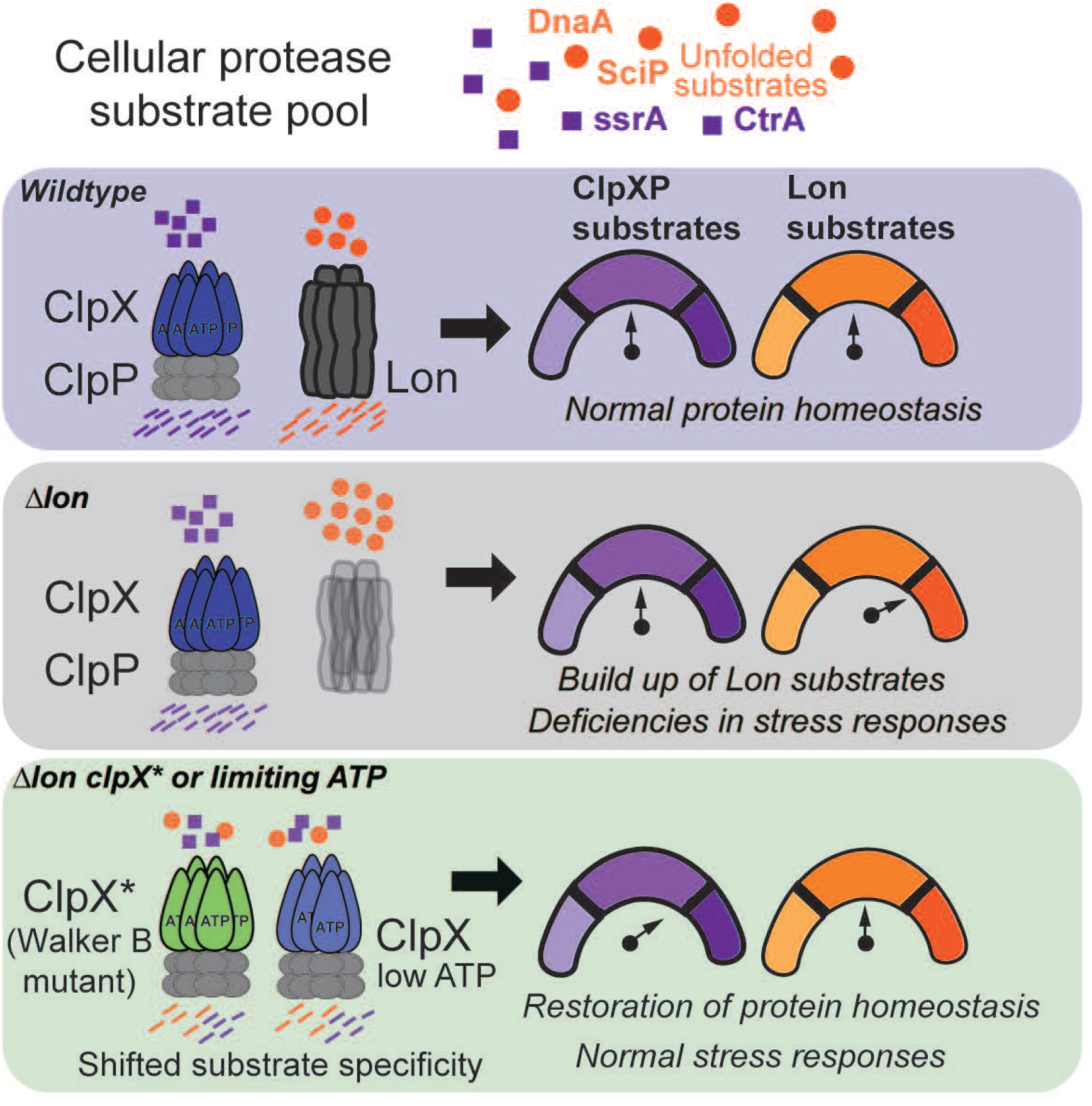

**eTOC:** AAA+ proteases, such as Lon and ClpXP, select distinct targets for degradation to maintain proteostasis. Mahmoud et al. show that ATP hydrolysis can tune substrate specificity of ClpX, allowing ClpX to degrade Lon-restricted substrates under limiting ATP conditions or in the presence of a ClpX mutant.

**Highlights:**

- A Walker B mutation of the AAA+ protease ClpX alters substrate specificity
- ClpX mutant degrades new substrates but degrades canonical substrates less well
- Decreasing ATP levels enhances ClpXP mediated degradation of some classes of substrates
- ATP-induced changes in conformational states accompany alterations in ClpX specificity

## Introduction

Energy-dependent protein degradation regulates normal growth and stress responses in all cells. In bacteria, regulated proteolysis is carried out by several energy-dependent AAA+ (ATPases associated with cellular activities) proteases which include Lon, ClpXP, ClpAP, HslUV, FtsH (Gur et al., 2011; Mahmoud and Chien, 2018). These proteases share a similar general architecture containing an unfoldase domain and a non-specific peptidase (Sauer and Baker, 2011). The two domains can be encoded on a single polypeptide, such as in the case of Lon and FtsH, or they can be encoded by separate proteins, an ATPase, such as ClpX, and a peptidase, such as ClpP (Baker and Sauer, 2012; Langklotz et al., 2012; Lee and Suzuki, 2008). Using the power of ATP hydrolysis, the unfoldase recognizes, unfolds, and translocates substrates into the sequestered peptidase chamber to be degraded. Because they are critical for maintaining proteostasis, loss of these proteases results in defects in growth, cell-cycle progression, stress responses, and pathogenesis (Breidenstein et al., 2012; Howard-Flanders et al., 1964; Jenal and Fuchs, 1998; Rogers et al., 2016).

Despite the similarities in both architecture and mechanism, these proteases generally have distinct niches of substrate preference. For example, in *Caulobacter crescentus*, Lon is the principal protease for degradation of the replication initiator DnaA (Jonas et al., 2013), the methyltransferase CcrM (Wright et al., 1996), and the transcriptional regulator SciP (Gora et al., 2013). By contrast, the ClpXP protease degrades cell-cycle factors like CtrA and TacA during cell-cycle progression through an adaptor hierarchy (Joshi et al., 2015). Lon is also responsible for misfolded protein degradation (Goldberg, 1972; Gur and Sauer, 2008; Jonas et al., 2013). However, while a handful of target substrates for these AAA+ proteases have been identified, it remains unclear how proteases are able to discriminate among protein targets and recognize distinct substrates for irreversible degradation.

In this work, we find that a variant of ClpX (*clpX*\*) can compensate for the absence of the Lon protease. Expression of *clpX*\* suppresses defects in motility, growth, filamentation, and sensitivity to stress normally seen in a Δ*lon* strain. In addition to this phenotypic rescue, degradation of normal Lon substrates is restored *in vivo* and *in vitro* by ClpX*P. This increased ability to degrade noncanonical substrates comes at the cost of reduced degradation of ClpXP-specific substrates, resulting in fitness defects in otherwise wildtype strains expressing the *clpX*\* allele. Further mechanistic characterization shows that ClpX* has reduced catalytic efficiency for ATP hydrolysis, suggesting a connection between ATP utilization and substrate specificity. Consistent with this, limiting ATP for wildtype ClpXP shifts substrate preferences similar to that of ClpX*P. Limited proteolysis, differential scanning fluorimetry (DSF), and hydrogen-deuterium exchange mass spectrometry (HDX-MS) show that the ClpX unfoldase adopts distinct conformations under these ATP limiting or saturating conditions. Taken together, our work demonstrates that ATP-dependent dynamics between conformational states are important for substrate recognition by AAA+ unfoldases.

## Results

### A suppressor screen identifies a *clpX* mutant which rescues *Δlon*

We reasoned that a suppressor screen would allow us to identify Lon-related interactions and used a transposon library to identify mutations that restore motility to a *Δlon* strain. We isolated a high motility mutant with a transposon insertion in CCNA_00264; however, transduction experiments revealed that this transposon alone was not able to rescue motility (Figure S1A). Sequencing the genome of the suppressor strain showed a point mutation in *clpX*, which resulted in a single amino acid change from glycine 178 to alanine (*clpX*\*). This glycine is highly conserved among ClpX homologs and immediately adjacent to the Walker B motif (Figure S1C). By generating the *clpX*\* allele in a *Δlon* background, we found that *clpX*\* on its own partially restores motility (Figure 1A).

**Figure 1:**
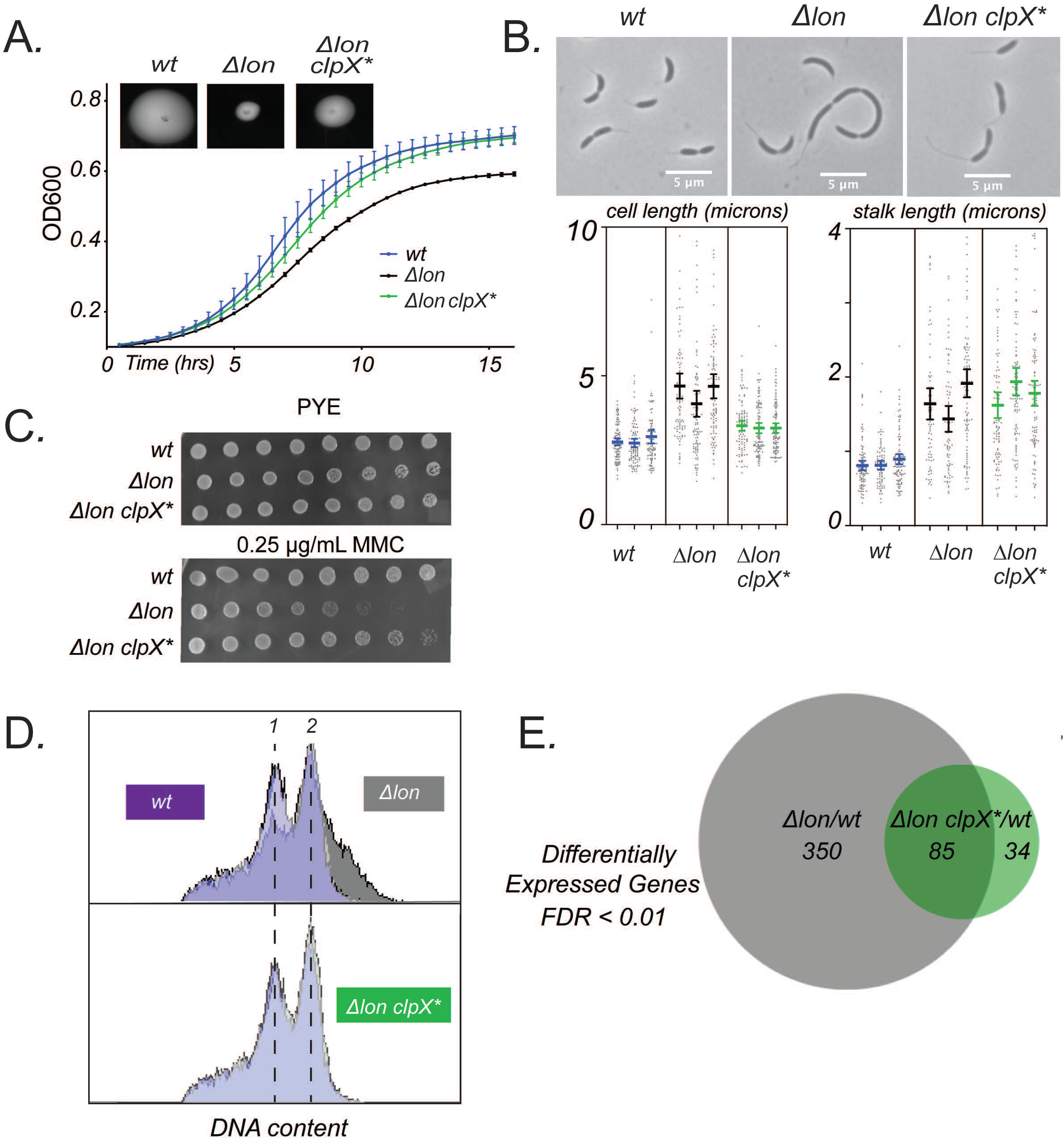
*clpX*\* mutant suppresses *Δlon* phenotypes. A. Growth curves of *wild type* (*wt*), *Δlon*, and *Δlon* clpX* cells grown in PYE. Biological triplicate experiments are shown. Error bars represent 95% confidence interval. Inset shows motility assays as measured by growth in 0.3% PYE agar. B. Representative phase-contrast microscopy images of *wt*, *Δlon*, and *Δlon* clpX* cells grown in PYE during exponential phase. Quantification of cell length and stalk length for three biological replicates of n=100 cells. C. Serial dilution assays comparing colony formation of strains in PYE and PYE supplemented with mitomycin C. Spots are plated 10-fold dilutions of exponentially growing cells from left to right. D. Flow cytometry profiles showing chromosome content of indicated strains after three-hour treatment with rifampicin. Cells were stained with SYTOX Green to measure DNA content. The fluorescent intensities corresponding to one chromosome (1) and two chromosomes (2) are indicated. Experiment was performed two times. Representative data from one of the biological replicates is shown here. E. Venn diagram summarizing the number of differentially expressed genes with an FDR cutoff < 0.01 from RNA seq performed with stationary phase cells. Venn diagram created by BioVenn (Hulsen, de Vlieg and Alkema, 2008). See also Figure S1 and Supplementary File 1.

We were intrigued that a single mutation in *clpX* could restore motility to cells lacking Lon and chose to further characterize this strain. Cells lacking Lon show growth defects that exhibit as an extended lag phase and reduced cell mass accumulation in stationary phase (Figure 1A). The *clpX*\* allele rescues the mass accumulation defect, but not the extended lag (Figure 1A). *Δlon* strains also have significant morphological deficiencies including elongated cell length and longer stalks, external polar organelles characteristic of *Caulobacter*. We found that the *clpX*\* mutant restored the cell length of Δ*lon* strains to near wildtype levels (Figure 1B, 1C). However, stalks remained elongated in the *Δlon clpX*\* strain (Figure 1B, 1C).

We next wondered if, in addition to suppression of *Δlon* morphological and growth defects, sensitivity to certain stressors would be suppressed by this *clpX*\* allele. One of the originally described phenotypes for *E. coli lon* mutants is DNA damage sensitivity (Mizusawa and Gottesman, 1983; Witkin, 1946) and *Caulobacter Δlon* strains are also highly sensitive to various DNA damaging agents (Zeinert et al., 2018), including mitomycin C (MMC) (Figure 1C). We found that *Δlon clpX*\* was 100-fold more resistant to MMC than *Δlon* alone (Figure 1C).

As mentioned previously, Lon is the major protease responsible for the degradation of the replication initiator DnaA and *Δlon* strains accumulate excess chromosome content (Jonas et al., 2013; Wright et al., 1996). Like the morphological abnormalities, over-replication is suppressed in the *Δlon clpX*\* strain which shows similar chromosome content as wildtype cells (Figure 1D, S1D). To determine if this rescue extended to a systems-level correction, we compared RNA-seq profiles of wildtype, *Δlon* and *Δlon clpX*\* strains. As expected, many genes (435) are differentially expressed upon loss of Lon (Figure 1E, Supplementary File 1), while the *Δlon clpX*\* strain shows fewer differences (119) with only 85 genes overlapping between these sets. Finally, we assayed the overall physiological consequence of ClpX* using competitive fitness in mixed cultures. We grew cultures of *Δlon* or the *Δlon clpX*\* strain together with *Δlon* strains constitutively expressing the Venus fluorescent protein (Persat et al., 2014). We found that *Δlon clpX*\* cells were more fit than *Δlon* cells (Figure S1E). Our interpretation is that expression of *clpX*\* in a *Δlon* background largely shifts the phenotypes and transcriptional landscape to more wildtype profiles.

### ClpX* restores degradation of Lon substrates *in vivo*

Because cells lacking Lon would accumulate normally degraded substrates, we hypothesized that the rescue we observed in the *Δlon clpX*\* strain was likely due to lower levels of Lon substrates, such as DnaA, CcrM, and SciP (Figure 2A) (Wright et al., 1996; Gora et al., 2013; Jonas et al., 2013). As expected, in unsynchronized cells, DnaA and CcrM levels are higher in the *Δlon* strain than in wildtype (Figure 2B). Similarly, SciP levels are higher in *Δlon* swarmer cells in comparison to wildtype (Figure 2D). Interestingly, levels of DnaA and SciP are restored to wildtype levels in the *Δlon clpX*\* strain; however, CcrM remained at higher levels (Figure 2B). The reduced levels of DnaA likely explain the rescue of replication defects and enrichment analysis of the RNA-seq results shows that the SciP-controlled regulon and cell-cycle gene expression are largely restored in the *Δlon clpX*\* strain (Figure S2). We reasoned that expression of *clpX*\* rescues normal growth and critical stress responses in a *Δlon* strain through restoring wildtype levels of key proteins.

**Figure 2:**
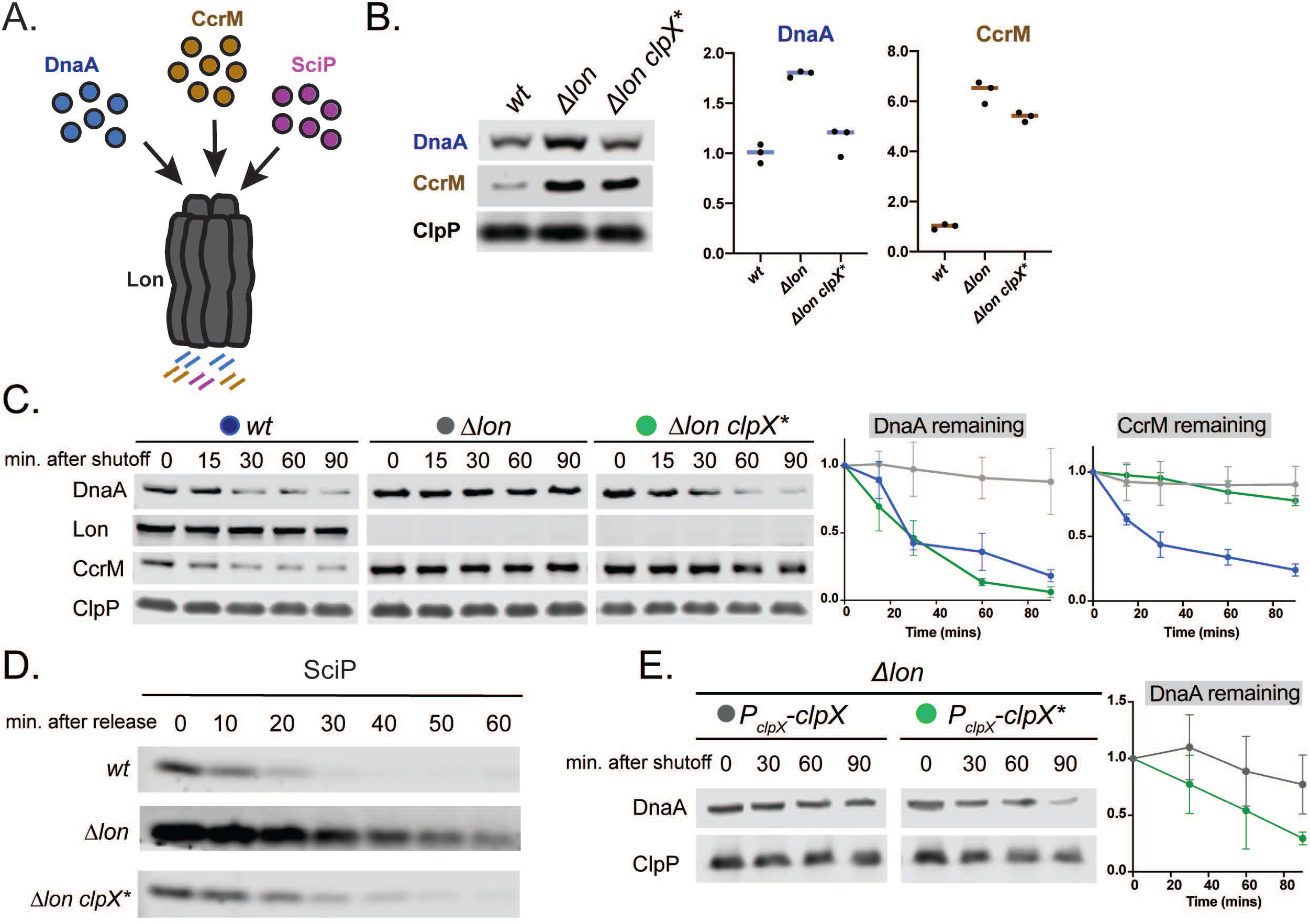
*clpX*\* mutant restores levels of Lon substrates through degradation. A. Lon degrades DnaA, SciP, and CcrM in *C. crescentus* (Wright *et al*., 1996; Gora *et al*., 2013; Jonas *et al*., 2013). B. Western blot showing DnaA and CcrM levels in *wt*, *Δlon*, and *Δlon* clpX* cells. Lysates from an equal number of exponential phase cells were probed with anti-DnaA or anti-CcrM antibody. ClpP was used as a loading control. A representative image and quantifications of triplicate experiments are shown. Line represents the mean. C. Antibiotic shutoff assays to monitor DnaA and CcrM stabilities in wt, *Δlon, and Δ*lon* clpX*\* cells. Chloramphenicol was added to stop synthesis and lysates from samples at the indicated time points were used for western blot analysis. Quantifications of triplicate experiments with substrate shown relative to ClpP control levels are shown to the right. Error bars represent SD. D. Western blot showing SciP levels in synchronized populations of wt, *Δlon*, and *Δlon clpX*\* cells. Swarmer cells were isolated using a density gradient and an equal number of cells were released into fresh PYE medium. Samples were withdrawn at the indicated time points and probed with anti-SciP. E. DnaA stability is measured in *Δlon* strains expressing an extra copy of wildtype *clpX* or *clpX*\*. Experiment was performed four times. Representative data from one of the biological replicates is shown here. Quantifications of experiments shown to the right. Error bars represent SD. See also Figure S2.

Next, we explored the mechanism by which a mutation in ClpX restores steady state levels of DnaA and SciP. Because DnaA and SciP are controlled by proteolysis, we suspected that protein turnover may be affected. As expected, DnaA is largely stabilized in *Δlon* strains (Figure 2C; Jonas, et al. 2013); however, we found that DnaA degradation was restored in the Δ*lon clpX* strain*, with a half-life similar to that of wildtype (Figure 2D). SciP protein dynamics and cell-cycle phase specific levels were also restored in *Δlon clpX*\* cells (Figure 2D). By contrast, CcrM degradation was still solely dependent on Lon, even when *clpX*\* was present, in both unsynchronized cells (Figure 2C) and during cell-cycle progression (Figure S2D).

We considered that changes in protein turnover could be either due to a gain-of-function in the *clpX*\* mutant or by indirect cellular effects associated with a loss-of-function in *clpX*. To test this, we created a merodiploid *Δlon* strain expressing a second copy of *clpX* or *clpX*\* under control of the native promoter along with a normal chromosomal copy of *clpX*. We reasoned that if the effect was direct, then expression of *clpX*\* would be sufficient to restore DnaA degradation. If the effect was indirect, then DnaA would remain stable. Consistent with a gain-of-function phenotype for the *clpX*\* allele, we found that DnaA turnover increased upon expression of *clpX*\* but not wildtype *clpX* (Figure 2E). Similarly, the phenotypic rescue we observed in the *Δlon clpX*\* strain was also due to a gain-of-function activity as the merodiploid strain expressing *clpX*\* suppressed *Δlon* defects, such as growth, motility, and genotoxic stress tolerance (Figure S2E-G). Because ClpXP is a AAA+ protease, we hypothesized that ClpX*P (the protease complex of the variant ClpX* with wildtype ClpP) was now able to recognize and degrade substrates normally degraded by Lon.

### ClpX*P directly degrades Lon substrates *in vitro*

As a direct test of our hypothesis, we purified the ClpX* variant and reconstituted degradation *in vitro*. Upon initial characterization of purified ClpX*, we found that it formed an active ATPase, but hydrolyzed ATP three times faster than wildtype (Figure S3A). Consistent with the fact that Lon is the principal protease for DnaA degradation (Jonas et al., 2013; Liu et al., 2016), we found that wildtype ClpXP poorly degraded DnaA *in vitro* (Figure 3A). However, purified ClpX*P could degrade DnaA four-fold faster than ClpXP (Figure 3A). Similarly, SciP was degraded three-fold faster by ClpX*P in comparison to ClpXP (Figure 3A). Finally, as predicted from our *in vivo* results, CcrM was not degraded by either ClpX*P or ClpXP (Figure 3A, Figure S3B), but was well degraded by Lon (Figure S3B). Thus, ClpX* appears to have an expanded substrate profile which includes some Lon substrates, but others, such as CcrM, remain exclusive to Lon.

**Figure 3:**
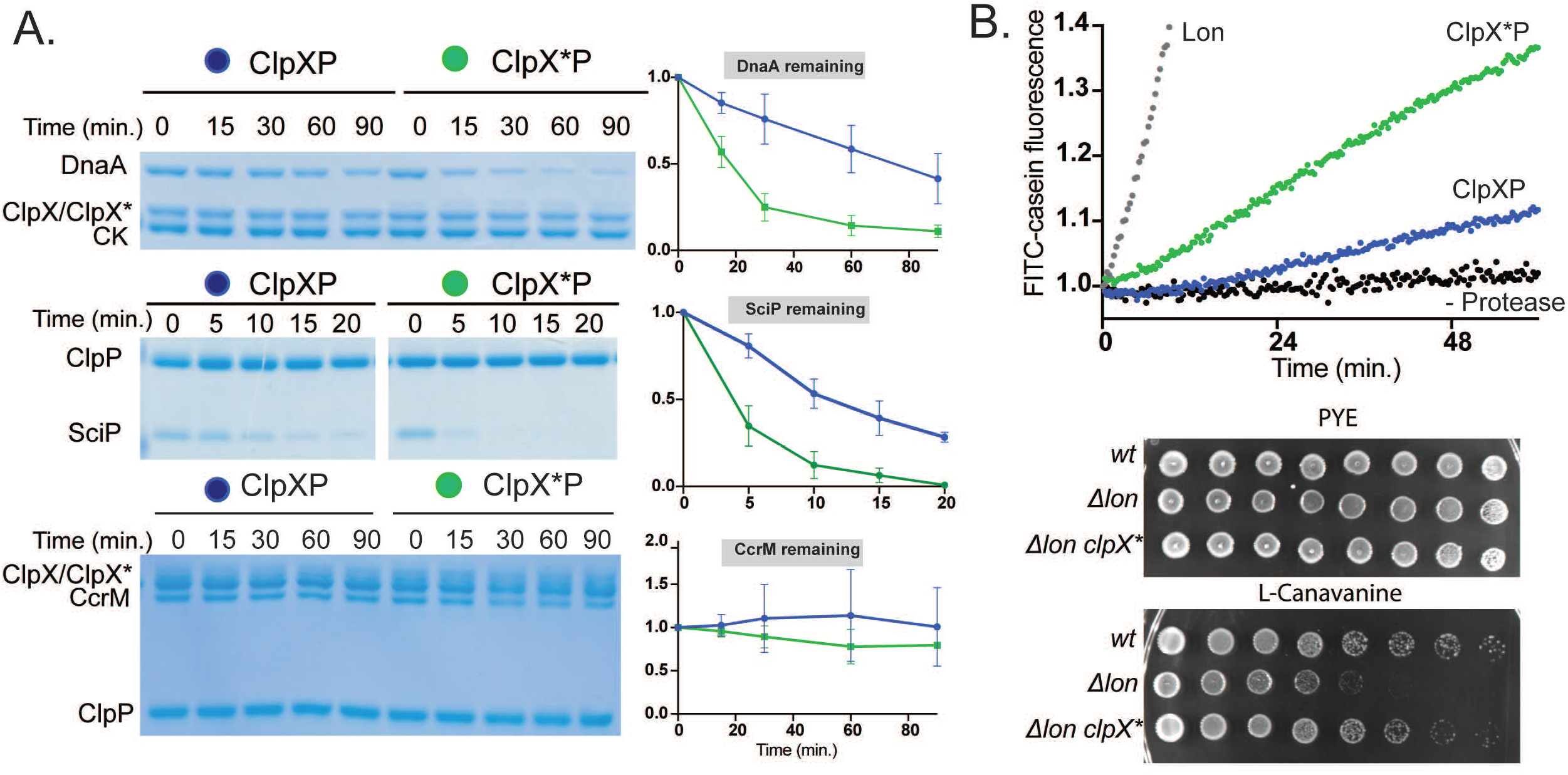
ClpX*P degrades some Lon substrates faster than ClpXP *in vitro*. A. *In vitro* degradation of DnaA, SciP, or CcrM. Assays were performed with 0.1 μM ClpX_6_ or ClpX_6_*, and 0.2 μM ClpP_14_. Substrate concentrations were 1 μM DnaA, 5 μM SciP, 0.5 μM CcrM. Quantification of triplicate experiments shown. Error bars represent SD. B. *In vitro* fluorescence degradation assay of FITC-casein in the presence of ClpXP or ClpX*P or Lon. Degradation assays were performed with 10μg/mL FITC-Casein, 0.1 μM ClpX_6_ or ClpX_6_* and 0.2 μM ClpP_14_ or 0.1 μM Lon_6_. Serial dilution assays comparing colony formation of strains in PYE and PYE supplemented with L-canavanine. Spots are plated 10-fold dilutions of exponentially growing cells from left to right. See also Figure S3.

Lon recognizes exposed hydrophobic amino acids (Gur and Sauer, 2008) and is best known as a quality control protease that eliminates mis/unfolded proteins (Goff et al., 1984). Given its altered specificity, we considered if ClpX* could now recognize these substrates. Casein is commonly used as an *in vitro* model substrate for mis/unfolded proteins. Lon robustly degrades fluorescently labeled casein, whereas wildtype ClpXP poorly degrades this substrate (Figure 3B). Intriguingly, ClpX*P degrades FITC-casein more than twice as fast as ClpXP (Figure 3B, Figure S3C). Similar to casein, ClpX*P degrades another unfolded Lon substrate, a chemically denatured domain of titin tagged with the hydrophobic sequence β20 (Gur and Sauer, 2008), better than wildtype ClpXP (Figure S3D).

Because cells lacking Lon are sensitive to proteotoxic stresses that create misfolded proteins and because ClpX*P is better at degrading mis/unfolded substrates than wildtype ClpXP, we wondered if the *Δlon clpX*\* strain would be more resistant to stressors that induce cellular protein misfolding. Indeed, the presence of the *clpX*\* allele protected *Δlon* cells from hypersensitivity to the proteotoxic amino acid L-canavanine (Figure 3B). We conclude that ClpX* has an altered substrate preference, which includes DnaA, SciP, and unfolded proteins, whose degradation are normally governed by the Lon protease.

### ClpX*P is deficient in degradation of native ClpXP substrates

Mutations in ClpX that improve recognition of some substrates can have negative consequences on others (Farrell et al., 2007). GFP-ssrA is a well-characterized ClpXP substrate that is directly bound by the pore loops of ClpX (Fei et al., 2020; Gottesman et al., 1998). We purified GFP fused to the *Caulobacter* ssrA tag (Chien, et al. 2007), determined degradation rates by ClpX*P and ClpXP, and fit these rates to the Michaelis-Menten equation. We found that ClpX*P degrades GFP-ssrA more poorly than wildtype ClpXP, principally lowering the turnover rate (Figure 4A, Figure S4A), suggesting that either ClpX* fails to irreversibly initiate degradation via the ssrA tag as readily or is unable to translocate the reporter protein as well as wildtype ClpX.

**Figure 4:**
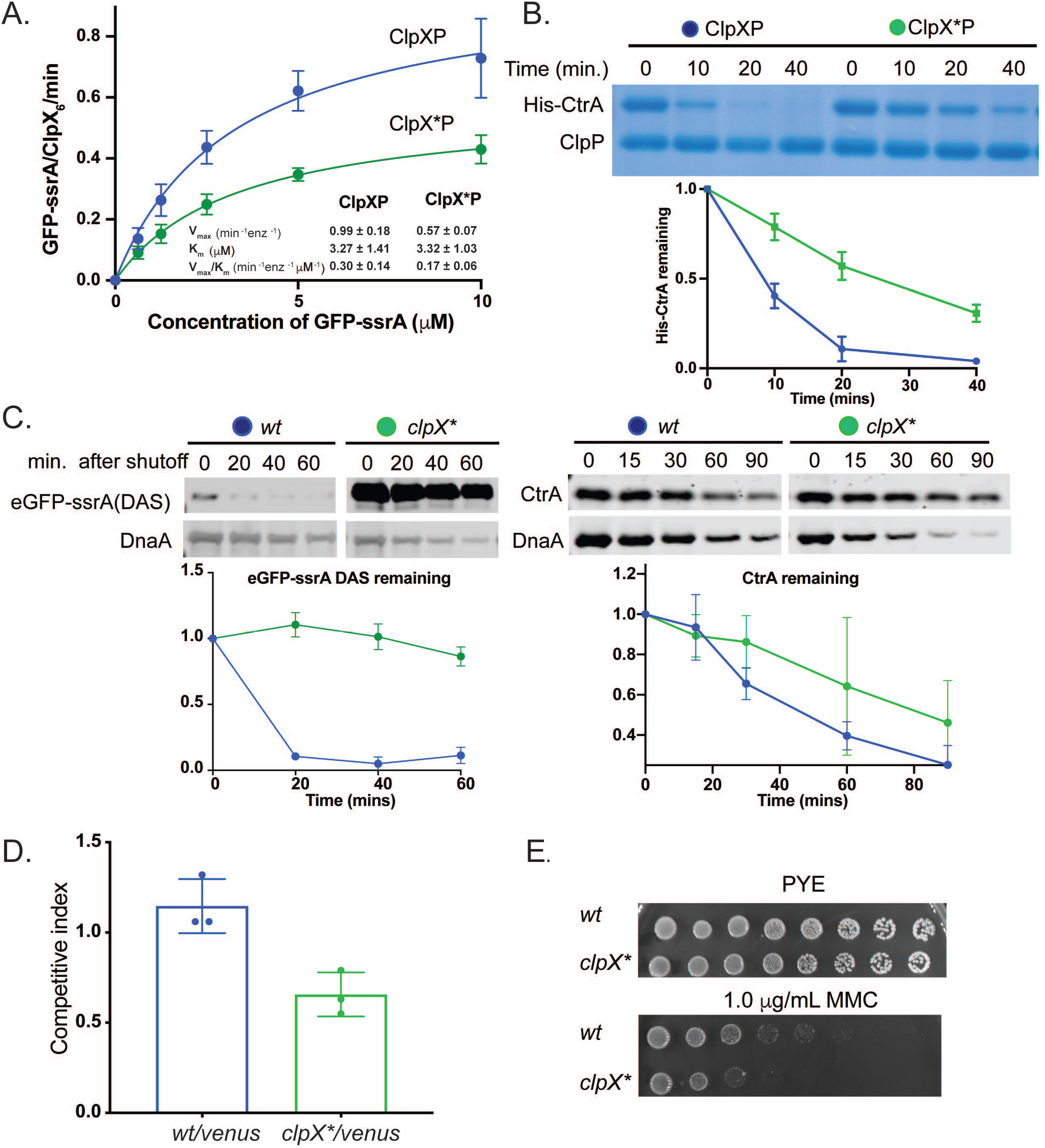
ClpX* mutant is deficient in degradation of native ClpXP substrates. A. Michaelis-Menten plot showing the rate of degradation as a function of GFP-ssrA concentration by ClpXP and ClpX*P. Inset displays kinetic parameters. Assays were performed with 0.1 μM ClpX_6_ or ClpX*_6_, 0.2 μM ClpP_14_, ATP regeneration system, and the indicated concentrations of GFP-ssrA. Data was fitted to Michaelis-Menten equation. Triplicate experiments are shown. B. *In vitro* degradation assay of His-CtrA in the presence of ClpXP or ClpX*P. Degradation assays were performed with 3 μM His-CtrA, 0.1 μM ClpX_6_ or ClpX*_6_, 0.2 μM ClpP_14_, and ATP regeneration system. Quantification of duplicate experiments is shown below. Error bars represent SD. C. *In vivo* degradation assay showing eGFP-ssrA (DAS), DnaA, and CtrA stability in *wt* and *clpX*\* cells. The plasmid encoded M2FLAG-eGFP-ssrA (DAS) construct was induced with the addition of 0.2% xylose. Chloramphenicol was used to inhibit protein synthesis. Samples were withdrawn at the indicated time points and quenched in SDS lysis buffer. Lysate from an equal number of cells was used for western blot analysis and probed with anti-M2, or anti-CtrA, and anti-DnaA antibody. Quantification of triplicate experiments is shown below. Error bars represent SD. D. Competition assay with wildtype cells harboring xylX::Plac-venus (constitutive venus expression) and nonfluorescent wt or *clpX*\* strains. exponential phase cells were mixed 1:1, diluted, and allowed to outgrow for 12 doublings. Quantification of triplicate experiments is shown below. Error bars represent SD. E. Serial dilution assays comparing colony formation of *wt* and *clpX*\* cells in PYE and PYE supplemented with MMC. Spots are plated 10-fold dilutions of exponentially growing cells from left to right. See also Figure S4.

CtrA is a master regulator in *Caulobacter* that inhibits replication initiation and regulates the transcription of many cell-cycle genes (Laub et al., 2002; Wortinger et al., 2000). *In vivo*, CtrA is degraded during the G1-S transition by ClpXP through the use of an adaptor hierarchy (Domian et al., 1999; Jenal and Fuchs, 1998; Joshi et al., 2015; Quon et al., 1996); however, *in vitro* CtrA can also be directly recognized by ClpXP as its C-terminus resembles the ssrA tag (Chien et al., 2007). Consistent with a reduced ability to recognize ClpXP substrates, ClpX*P degrades isolated CtrA more poorly than wildtype (Figure 4B) and also shows reduced degradation of a CtrA-derived GFP reporter (Smith et al., 2014) in the presence of the full adaptor hierarchy (Figure S4B). We conclude that ClpX*P overall has a diminished ability to recognize and degrade native substrates normally degraded by ClpXP.

### Deficiencies of ClpX* revealed when Lon is present

The prior *in vitro* data suggests that rather than just expanding specificity, this mutation has allowed ClpX*P to degrade Lon-dependent substrates better but at the cost of degrading ClpXP-dependent ones. If this is true, while *clpX*\* is beneficial in a *Δlon* background, we reasoned that it may be detrimental to *lon^+^* cells.

To test this hypothesis, we generated an otherwise wildtype strain with *clpX*\* as its sole copy. We first examined ssrA tag turnover in this strain by using an ssrA derivative that relies on the SspB adaptor for degradation (eGFP-ssrA(DAS)) (Chowdhury et al., 2010) so that degradation would be sufficiently slowed to be visualized by western blotting. Consistent with our *in vitro* experiments, we observed stabilization and dramatically higher levels of eGFP-ssrA (DAS) in the *clpX*\* strain in comparison to wildtype (Figure 4C). We observed reduced CtrA degradation in the *clpX*\* strain, consistent with our *in vitro* observations (Figure 4C). Finally, DnaA degradation is accelerated in the *clpX*\* strains compared to wildtype – consistent with our findings that both Lon and ClpX*P are able to degrade DnaA (Figures 4C).

To assess the overall impact of ClpX* on wildtype cells, we performed competition assays as described previously using a wildtype strain constitutively expressing the Venus fluorescent protein (Persat et al., 2014). We found that while nonfluorescent wildtype cells could slightly outcompete the constitutive Venus expressing cells, the *clpX*\* strain showed a substantial competitive disadvantage (Figure 4D). We also tested for stress tolerances and found that *clpX*\* strains fail to survive genotoxic stress as robustly as wildtype (Figure 4E). Our conclusion is that reduced fitness of the wildtype *clpX*\* strain is likely due to reduced degradation of normal ClpXP substrates (such as CtrA) and prolific degradation of normal Lon substrates (such as DnaA) that together reduce overall fitness. Collectively, this supports our understanding that AAA+ proteases have been optimized for specific priorities in a cell – expansion of their substrate profile, as seen with *clpX*\*, can compensate for loss of other proteases, but at the cost of their ability to recognize their normal substrates.

### Limiting ATP alters wildtype ClpXP substrate specificity

To understand how this mutation in ClpX leads to enhanced degradation of certain substrates, we further characterized the ATPase activity of ClpX*. While ClpX* hydrolyzes ATP three-fold faster than wildtype ClpX at saturating ATP concentrations (Figure S4A), Michaelis-Menten experiments showed that the K_M_ increases by 5-6 fold and the k_cat_ for ATP hydrolysis increases by approximately 3 fold (Figure 5A). While K_M_ does depend on k_cat_, the magnitude of K_M_ increase given the change in k_cat_ suggests that there may also be deficiencies in nucleotide binding of ClpX*. To test this, we used ATP to compete off mant-ADP, a fluorescent ADP analog and we observed at least a three-fold increase in the IC_50_ for ClpX*, suggesting that it binds ATP worse than wildtype ClpX (Figure S5A). To make sure that the observed difference in the IC_50_ between ClpX and ClpX* was due to changes in nucleotide binding and not because of the hydrolysis of ATP in the assay, we also competed mant-ADP using ATPγS and assayed changes using fluorescence polarization. Consistent with the ATP titration described above, we calculated a higher IC_50_ for ATPγS for ClpX* (Figure S5B).

**Figure 5:**
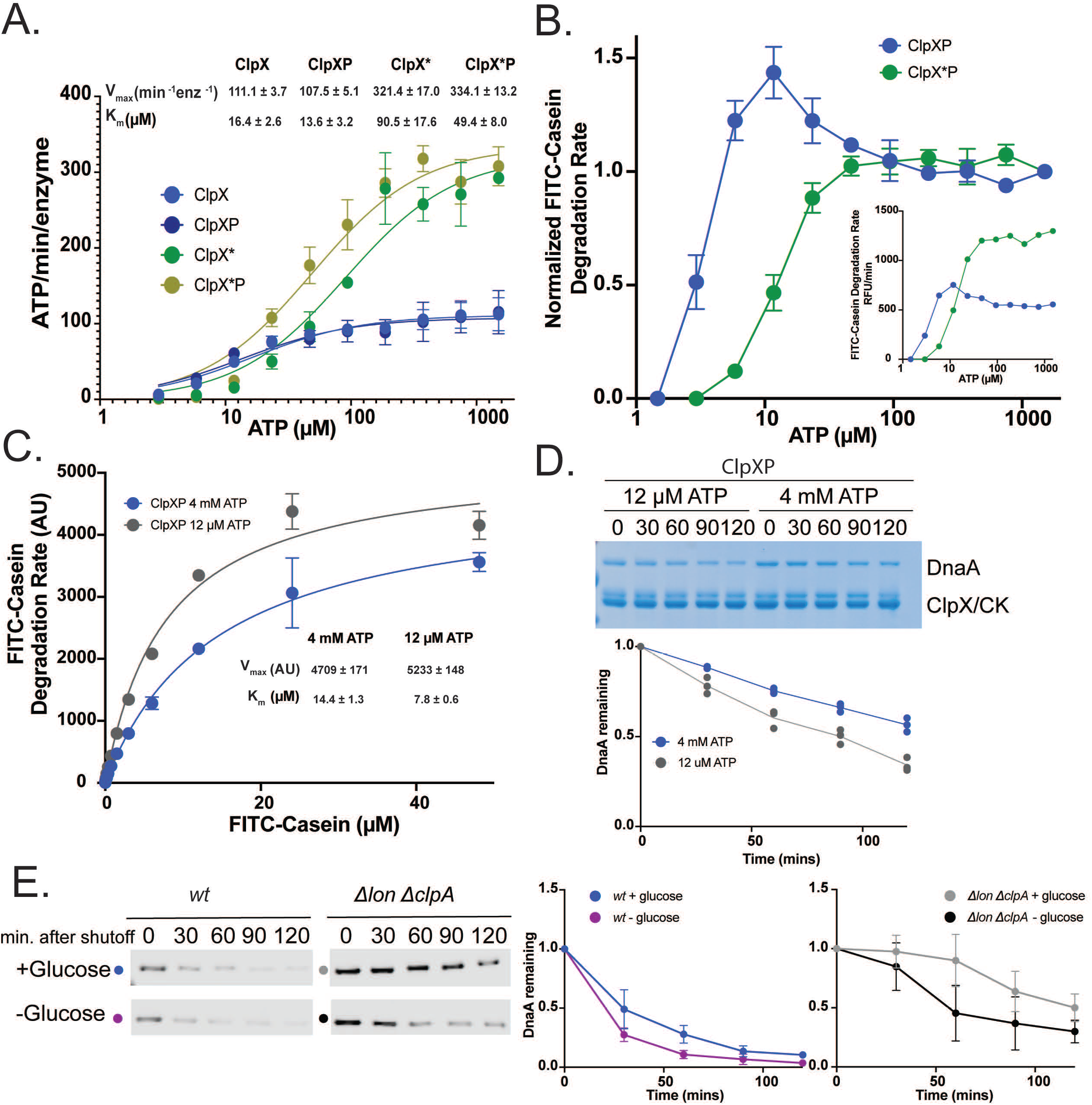
Limiting ATP alters wildtype ClpXP substrate specificity. A. Michaelis-Menten plot showing the rate of ClpX/ClpX* catalyzed ATP hydrolysis as a function of ATP concentration. Inset displays kinetic parameters. Assays were performed with 0.1 μM ClpX_6_ or ClpX*_6_, in the presence and absence of 0.2 μM ClpP_14_. Data was fitted to Michaelis-Menten equation. Triplicate experiments are shown. B. FITC-Casein degradation by ClpXP or ClpX*P as a function of ATP concentration. Assays were performed with 0.1 μM ClpX_6_ or ClpX*_6_, 0.2 μM ClpP_14_, ATP regeneration system, and 50 μg/mL FITC-Casein. Rates were normalized to highest concentration of ATP. Non-normalized rates shown in the inset. C. Michaelis-Menten plot showing the rate of degradation as a function of FITC-Casein concentration by ClpXP under low and saturating ATP conditions. Inset displays kinetic parameters. Assays were performed with 0.1 μM ClpX_6_, 0.2 μM ClpP_14_, and ATP regeneration system. Data was fitted to Michaelis-Menten equation. Triplicate experiments are shown. D. *In vitro* degradation of DnaA by ClpXP under low and saturating ATP conditions. Assays were performed with 0.1 μM ClpX_6_, and 0.2 μM ClpP_14_ and 0.5 μM DnaA. Quantification of triplicate experiments. E. Antibiotic shutoff assays to monitor DnaA under ATP limiting conditions in *wt* and *Δlon ΔclpA* strains. Chloramphenicol was added to stop synthesis and lysates from samples at the indicated time points were used for western blot analysis. Quantifications of triplicate experiments shown to the right. Error bars represent SD. See also Figure S5.

As seen previously, degradation of GFP-ssrA by ClpXP is precipitously lost below a minimum threshold of ATP (Martin et al., 2008), with the ClpX*P threshold being higher than that of wildtype, consistent with the higher K_M_ for ATP described above (Figure S5C). For ClpX*P, casein degradation reduced monotonically as ATP concentration was lowered (Figure 5B). In striking contrast, we found that for wildtype ClpXP, casein degradation was increased at intermediate nucleotide concentrations compared to saturating ATP (Figure 5B). Kinetic analysis shows that at these lower ATP concentrations ClpXP degrades casein with a lower K_M_ and a similar V_max_ compared to saturating ATP conditions (Figure 5C). Since K_M_ = K_D_ + k_cat_/k_on_, the lower K_M_ suggests that, at a minimum, casein must bind better to ClpX at these concentrations.

These results suggest that ClpX specificity depends on ATP concentration and rates of hydrolysis, with lower levels of ATP facilitating ClpX recognition of some substrates better than at saturating ATP concentrations. Similar to casein, we saw this same dependence *in vitro* with the Lon substrates DnaA and SciP (Figure 5D and S5D). To test this effect *in vivo*, we monitored degradation of DnaA during starvation, which should decrease cellular ATP levels, using a Δ*lon* Δ*clpA* strain to eliminate any non-ClpX dependent degradation (Liu et al., 2016). We verified that glucose starvation results in DnaA degradation in wildtype cells (Gorbatyuk and Marczynski, 2005; Figure 5E) and showed that starving *Δlon ΔclpA* also markedly increased DnaA degradation (Figure 5E). During this starvation there was also a drop in bulk ATP levels (Figure S5E), though we note that the magnitude of the drop is less than the changes we observe *in vitro* that alter ClpX specificity. We conclude that ClpX preference depends on ATP binding or hydrolysis, with its specificity expanded under limiting ATP conditions.

We considered if conformational changes in ClpX might explain the altered specificity of wildtype ClpX at low ATP or the changes associated with ClpX*. Using limited proteolysis to probe conformational differences we found that ClpX at low ATP was more protease accessible than at high ATP (Figure 6A). Similarly, protease accessibility of ClpX* at high ATP resembled the wildtype ClpX at low ATP (Figure 6A). Using differential scanning fluorimetry (DSF), we found that ClpX* was destabilized in comparison to wildtype ClpX at high ATP (Figure 6B). As expected, limiting ATP for wildtype ClpX also resulted in less thermal stability (Figure 6B) as ATP binding is known to be needed for ClpX oligomerization (Hersch *et al*., 2005). Taken together, these results suggest that under high ATP concentrations, the mutant ClpX* adopts a set of conformational states that is overall less stable that wildtype.

**Figure 6:**
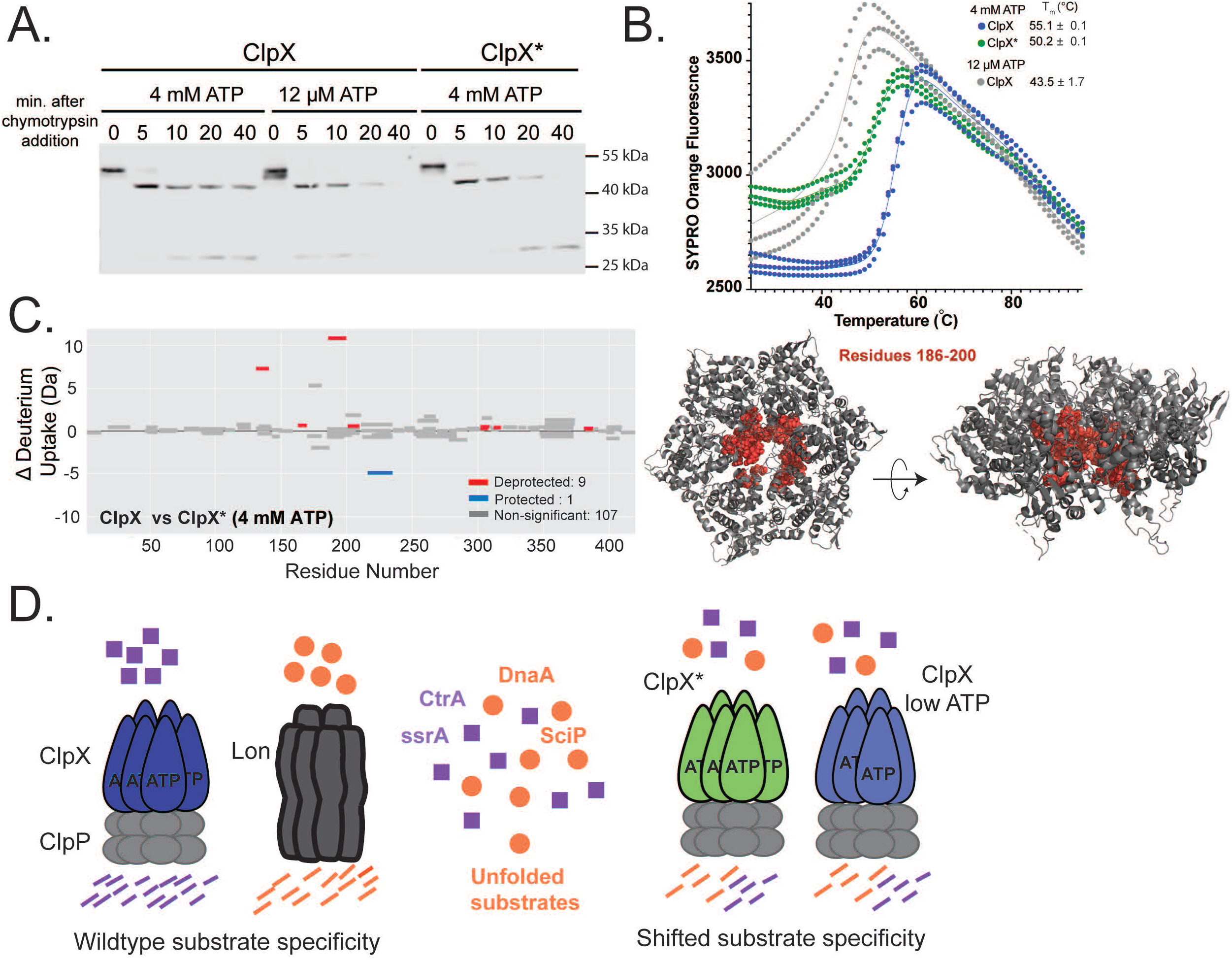
ClpX adopts distinct ATP-dependent conformational states. A. Limited proteolysis assay to probe conformational states. ClpX was incubated with Chymotrypsin with either 12 μM ATP or 4 mM ATP in the presence of ClpP and ATP regeneration system at 25 °C for the indicated time points. ClpX* was incubated with Chymotrypsin at 4 mM ATP in the presence of ClpP and ATP regeneration system at 25 °C for the indicated time points. ClpX was detected using anti-ClpX antibodies. B. Stability of ClpX* with 4 mM ATP, ClpX with 4 mM ATP, and ClpX with 12 μM ATP as measured by differential scanning fluorimetry. Triplicate experiments shown. C. Woods plot comparing deuterium uptake for wildtype ClpX vs ClpX* at 4 mM ATP after 60 minutes. Each bar on Woods plot represents a single peptide with peptide length corresponding to the bar length. Red bars indicate a deprotected (more deuterium uptake) region, blue represents a protected region, and gray bars are not significantly different. Woods plots were created with Deuteros (Lau et al. 2021) using the peptide significance test (p-value <0.01). Residues 186-200 are highlighted in red for the equivalent residues in the substrate-bound *E. coli* ClpX structure (Protein Data Bank ID: 6PO1) using PYMOL (Schrodinger). D. In a wildtype cell, AAA+ proteases Lon and ClpXP promote normal growth by degrading distinct substrates. ClpX*P can compensate for the absence of the Lon protease by tuning ClpX substrate specificity to better degrade Lon-restricted substrates (such as DnaA, SciP, and misfolded proteins) but this comes at the cost of native ClpXP substrates (such as ssrA-tagged proteins and CtrA). In ATP-limited conditions, wildtype ClpXP can undergo a similar switch in substrate specificity to ClpX*P, gaining the ability to better degrade DnaA, SciP, and misfolded proteins than the wildtype enzyme at saturating ATP. See also Figure S6.

To gain more clarity on what structural features might be different in these states, we performed hydrogen-deuterium exchange coupled with mass spectrometry (HDX-MS) to determine specific regions of ClpX and ClpX* that differ in dynamics. We compared ClpX and ClpX* in high ATP conditions (4 mM) and found that ClpX* peptides generally showed faster exchange than the equivalent peptides from wildtype ClpX (Figure 6C, S6B-C). One of these peptides (residues 186-200) was the most deprotected in ClpX* compared to ClpX and is buried in the pore of the closed substrate-bound structure of ClpX (Protein Data Bank ID: 6PO1) (Figure 6C, Figure S6C). Speculation on these results is discussed below.

## Discussion

Because proteolysis is irreversible, degradation of substrates must be carefully monitored to avoid toxic consequences. In bacteria, different energy dependent proteases have distinct substrate preferences that allows them to collectively regulate proteome dynamics. However, how these energy-dependent proteases choose their target substrates from a pool of cellular proteins remains poorly understood. Certain determinants that dictate specificity include degrons, or degradation tags, such as the ssrA tag which marks proteins for degradation by ClpXP or ClpAP (Keiler, 2015; Keiler et al., 1996). Specificity can be augmented through adaptor proteins that help deliver substrates to proteases. For example, degradation of CtrA in *Caulobacter* is restricted to the ClpXP protease via a highly regulated series of adaptors (Joshi et al., 2015).

In this work, we identified a mutant allele of *clpX* (*clpX*\*) that compensates for the loss of Lon through expanding ClpXP substrate specificity (Figure 6D). The variant ClpX*P protease complements the growth, motility, replication status, and morphology defects of a Δ*lon* strain (Figure 1A-1C) by restoring levels and degradation of the Lon substrates DnaA and SciP (Figure 2), which we confirm with biochemical experiments (Figure 3). Interestingly, ClpX*P has an improved ability to degrade unfolded protein substrates, a feature that likely explains the increased proteotoxic tolerance of *Δlon clpX*\* strains (Figure 3). Finally, this *clpX* variant is deficient in proteolysis of normal ClpXP substrates, such as CtrA and ssrA-tagged proteins, suggesting a shift in substrate preference rather than just an expansion (Figure 4, 6D). Mechanistic enzymology revealed that ClpX* requires ~5-fold more ATP for saturation of ATP hydrolysis (Figure 5A), leading us to explore the role of ATP levels in controlling specificity. Surprisingly, our *in vitro* studies found that wildtype ClpXP is better at degrading unfolded proteins, DnaA, and SciP in ATP limiting conditions, but worse at degrading ssrA-tagged proteins, similar to the behavior of ClpX*P even in saturating ATP conditions (Figure 5B, 5D, 6D, S5D). *In vivo*, DnaA degradation by wildtype ClpXP is accelerated (Figure 5E) in starvation conditions that also deplete ATP. We speculate that the features which alter ClpX* specificity even in high ATP conditions may also underlie the same change in specificity seen with wildtype ClpX under low ATP conditions.

How does altered ATP levels or altered ability to use ATP change ClpX substrate recognition? One model builds on emerging cryo-EM structures which have revealed shared features of substrate-bound AAA+ complexes, namely a shallow right-handed ring with pore loops gripping the substrate and at least 4 ATP-bound protomers (Fei et al., 2020; Gates et al., 2017; Lopez et al., 2020; Ripstein et al., 2020; Shin et al., 2020). Substrate-free forms of these machines show more open, often left-handed, spiral structures (Shin et al., 2020; Yokom et al., 2016). Our limited proteolysis experiments, which show faster chymotrypsin cleavage kinetics for ClpX* and wildtype ClpX at low ATP, suggest that they adopt distinct conformations from wildtype ClpX at saturating ATP (Figure 6A). Using HDX-MS, we find regions of ClpX* that show faster exchange than wildtype ClpX at saturating ATP (Figure 6C, Figure S6A).

One possible model that arrives from these results is that ClpX normally samples an ‘open’ state and a ‘closed’ state dependent on ATP levels. The‘closed’ state is the right-handed structure found in previous ClpXP structures (Fei et al., 2020; Ripstein et al., 2020), while the ‘open’ state would be more similar to the left-handed spiral structures found in substrate-free complexes (Shin et al., 2020; Yokom et al., 2016). We modeled this open state for ClpX by aligning six ClpX monomers with the AAA domains of substrate-free Lon from *Yersinia pestis* (Protein Data Bank ID: 6V11; Shin et al., 2020). The peptide corresponding to residues 186-200 was no longer buried in this modeled structure, consistent with a model where ClpX* can more readily adopt an open state (Figure S6D). In this case, limiting ATP or appropriate mutations can shift the open/closed balance toward the open state, which allows ClpX to better capture certain substrates, such as unfolded proteins, that may not have as strict sequence requirements for recognition (Figure S6E). However, degrons such as the ssrA tag which bind selectively to the closed conformation (Hersch et al., 2005) would be poorly recognized in these conditions (Figure S6E).

Alternatively, the changes in substrate specificity for ClpX* in saturating ATP conditions and wildtype ClpX under low ATP conditions could be due to entirely distinct mechanisms. For example, under low ATP concentrations, the ClpX oligomer is likely unstable. As evidence of this, our DSF experiments show additional destabilization of ClpX in low ATP conditions compared to either ClpX* and ClpX at saturating ATP (Figure 6B). Similarly, using HDX-MS, we observed that ClpX in low ATP conditions shows faster exchange across the entire sequence of ClpX in comparison to ClpX at high ATP (Figure S6A). The destabilization of the ClpX hexamer could be underlying the decreased GFP-ssrA degradation observed upon limiting ATP. However, this model does not explain why ClpX under limited ATP conditions degrades FITC-Casein, DnaA, and SciP better than at saturating ATP. Another possibility could be that ATP directly competes with these substrates for degradation. In this case, intermediate nucleotide concentrations provide enough ATP to power degradation and to also relieve inhibition, allowing ClpX to degrade these substrates. However, to date, there is no evidence that nucleotides directly compete with protein substrates for ClpX.

Prior work has shown that limiting nucleotide leads to loss of substrate degradation by ClpXP (Martin et al., 2008), as would make sense for an ATP-fueled unfoldase. We note that these studies exclusively used ssrA-tagged substrates, which require an ATP-bound (‘closed’) form of ClpX for recognition (Martin et al., 2008). Thus, the equivalent effects we are seeing here for casein, DnaA, and SciP would not have been previously observed, which highlights the importance of using a range of substrates in understanding activity. In our current work, we speculate that different states of ClpX, favored by either limiting ATP or mutation, can result in meaningful biological consequences. Intriguingly, similar conclusions were drawn for the chaperonin GroEL, where mutants with a shifted specificity that improves folding of one class of clients at the expense of others also resulted in an altered ATPase cycle attributed to changes in conformational lifetimes (Wang et al., 2002). Overall, it seems that exclusive substrate profiles for AAA+ systems are not hard-wired but can be altered not only by mutations but also through modulation of ATP hydrolysis, demonstrating a plasticity in these machines that yield flexibility in maintaining cellular proteostasis.

### Limitations of the study

While our final model that ATP affects the dynamics of open/closed states of ClpX oligomers satisfies our biochemical and cellular results, we acknowledge that there are limitations in the experimental support of our model. One of the major concerns is that ClpX oligomerization is ATP dependent, therefore under reduced ATP conditions the partial dissociation of ClpX into inactive monomers is a confounding factor in our interpretation of the results. A second concern is that although we favor a model where ClpX* at saturating ATP mimics wildtype ClpX under limiting ATP in terms of the mechanisms leading to shifted substrate specificity, it is possible that ClpX* has shifted substrate preference for a reason completely different than why wildtype ClpX under limiting ATP conditions has a similar shifted specificity. Because we have not directly measured ATP stoichiometry, we also cannot say for certain whether ClpX or ClpX* differ in nucleotide occupancy at saturating ATP concentrations. Finally, although open apo-state spirals and closed substrate-bound rings have been found for several AAA+ family members (as described above), these have yet to be directly seen for ClpX.

## Supporting information

Supplementary File 1: RNA Seq Analysis

Supplementary File 2: HDX Log File

## Acknowledgements

We thank the Chien, Strieter, Stratton, and Serio lab members for helpful comments and discussion on the manuscript. S.A.M. and B.A. were supported in part through the UMass NIH Chemistry Biology Interface Training Program (NIH T32GM008515), S.A.M. in part by the HHMI Gilliam Fellowship program, and B.A. in part by NIH R01GM135931. Work in the Chien lab is funded by the NIH (R35GM130320). We thank the Genomics Resource Laboratory (Institute for Applied Life Sciences; University of Massachusetts Amherst) and the University of Massachusetts Mass Spectrometry Core Facilities (RRID:SCR_019063) for services and equipment support.

## Author Contributions

Conceptualization: S.A.M. and P.C.; Investigation, S.A.M. and B.A.; Writing, S.A.M and P.C.; Funding Acquisition, P.C. Supervision, P.C.

## Declaration of Interests

The authors declare no competing interests

## STAR Methods

### RESOURCE AVAILABILITY

#### Lead Contact

- Further information and requests for resources and reagents should be directed to and will be fulfilled by the Lead Contact, Peter Chien.

#### Materials Availability

- All unique/stable reagents generated in this study are available from the Lead Contact without restriction.

#### Data and Code Availability

- RNAseq data are available in the NCBI Gene Expression Omnibus (GEO): GSE166765 (https://www.ncbi.nlm.nih.gov/geo/query/acc.cgi?acc=GSE166765)
- This paper does not report original code.
- Any additional information required to reanalyze the data reported in this paper is available from the lead contact upon request.

### EXPERIMENTAL MODEL AND SUBJECT DETAILS

#### Bacterial strains

Bacterial strains used in this study are listed in the key resources table. *Caulobacter crescentus* strains were grown in PYE medium (2g/L peptone, 1g/L yeast extract, 1 mM MgSO_4_, and 0.5 mM CaCl_2_) at 30°C and supplemented with antibiotics at the following concentrations: kanamycin (25 μg/ml), spectinomycin (100 μg/ml), and oxytetracycline (2 μg/ml). After initial selection steps for strain construction (excluding plasmids), antibiotics were excluded for all assays.

*Escherichia coli* strains were grown in LB (10g/L NaCl, 10g/L tryptone, 5g/L yeast extract) and supplemented with antibiotics at the following concentrations: ampicillin (100 ug/ml), oxytetracycline (15 μg/ml), spectinomycin (50 μg/ml), kanamycin (50 μg/ml), gentamycin (20 μg/ml). For solid medium, 1.5% agar was used regardless of bacterial species. For all strains, optical density was measured at 600 nm.

For induction purposes, the 477 plasmids were induced with xylose (0.2%) and repressed with glucose (0.2%). For protein expression, 0.4 mM IPTG was used for induction.

### METHOD DETAILS

#### Cloning and strain constructions

All *Caulobacter* strains were derived from CPC176, an isolate of NA1000. To generate the *Δlon clpX*\* strain, a two-step recombination protocol with a sucrose counter selection was utilized (Skerker et al., 2005). The vector pNPTS138 was digested with HindIII and EcoRI. A PCR product of the *clpX*\* sequence was amplified from the original motility suppressor and Gibson assembly was then used to generate pNPTS138_clpX*. Following transformation with pNPTS138_clpX* into NA1000 or *Δlon*, primary selection was on PYE supplemented with kanamycin. Primary colonies were grown overnight without selection and overnight cultures were played on PYE agar supplemented with 3% w/v sucrose. To validate allelic swap, strains were tested for sensitivity to kanamycin. *Δlon clpX*\* was also screened by motility and sequencing of the *clpX* locus validated candidate clones.

*ClpX* or *clpX*\* merodiploid strains were generated by Gibson assembly of PCR product and double digested plasmids of pMCS-2. The plasmids were then electroporated into *Δlon* and selected onto kanamycin plates. Strains were validated by anti-ClpX westerns.

eGFP-ssrA(DAS) induction strains were generated by electroporating eGFP-ssrA(DAS) 477 plasmid into wildtype and wildtype *clpX*\* and selecting on spectinomycin. Strains were validated by anti-M2 westerns.

#### Motility suppressor screen

Transposon libraries were generated for *Δlon*. Two-liter PYE cultures were grown to mid exponential phase, pelleted, and washed with 10% glycerol. Competent cells were electroporated with Ez-Tn5 <Kan-2> transposome (Lucigen, Madison, WI). Cells recovered for 90 minutes at 30 °C and then played on PYE plates supplemented with kanamycin. Libraries were grown for 7 days. Colonies were then scraped from the surface, combined, and resuspended to form a homogenous solution of PYE + 20% glycerol.

The Tn library was thawed out and diluted into a flask containing two-liter 0.3% agar. The cell agar mixture was plated and grown at 30 °C for 3 to 5 days. Candidates that appeared motile were validated by inoculating single colonies into motility agar on the same plate as NA1000 as a positive control and Δ*lon* as a negative control and incubating plates for 2-3 days at 30 °C.

#### Motility assay

A single colony was inoculated into PYE with 0.3% agar using a sterile pipette tip and left to incubate at 30°C for 2 to 3 days.

#### Whole genome sequencing

Genomic DNA was extracted using the MasterPure Complete DNA and RNA purification kit (Epicenter Biotechnologies, Madison, WI). A Qubit Fluorometer (ThermoFisher Scientific, Waltham, MA) was utilized to assess DNA concentration. Illumina libraries were generated from the extracted genomic DNA using the NexteraXT (Illumina, San Diego, CA) protocol. Libraries were multiplexed and sequenced at the University of Massachusetts Amherst Genomics Core Facility on the NextSeq 500 (Illumina). Single nucleotide polymorphisms (SNPs) were detected using breseq (Deatherage and Barrick, 2014).

#### RNA sequencing

RNA was extracted from stationary phase cells. Libraries were generated from the extracted RNA using the NEB Next RNA Library Prep kits (NEB, Ipswich, MA). Libraries were sequenced at the University of Massachusetts Genomics Core Facility on the NextSeq 500 with single end 75 base reads.

#### Plating viability and drug sensitivity

All *Caulobacter* strains were grown overnight in liquid PYE media. After overnight growth, cells were back diluted to OD600 0.1 and outgrown to mid-exponential phase before being normalized to OD_600_ 0.1 and 10-fold serially diluted on to media. For experiments using mitomycin C (Sigma, St. Louis, MO) and L-canavanine (Sigma), drugs were prepared at a stock concentration of 400 μg/ml mitomycin C, and 100 mg/ml L-canavanine and filter sterilized. PYE agar was cooled before the drugs were added and plates were left to air dry prior to serial dilution plating. All plates were incubated at 30 °C for 2-3 days and imaged with a Syngene G:Box.

#### *In vivo* assays

The stability of proteins *in vivo* was determined by inhibiting protein synthesis upon addition of 30 μg/ml chloramphenicol to cells in exponential phase. At each time point, 1ml of culture was removed and centrifuged at 15,000 rpm for 2 minutes. The supernatant was removed and pellets were flash frozen in liquid nitrogen. Pellets were thawed, resuspended in 2× SDS dye, and normalized to the OD_600_ of the lowest sample. Samples were boiled for 10 minutes and centrifuged for 10 minutes at 15,000 rpm. Extracts were run on 10% Bis-Tris gels for 1 hour at room temperature at 150 V. Gels were then transferred to nitrocellulose membranes for 1 hour at room temperature at 20V. Membranes were blocked with 3% milk in Tris-based saline with 0.05% Tween-20 (TBST) for 1 hour. Membranes were probed with primary antibody in 3% milk in TBST at 4°C overnight with following dilution factors: 1:5000 dilution of DnaA, 1:5000 SciP, 1:5000 CcrM, 1:5000 ClpP, 1:2500 Lon, and 1:2000 M2. Membranes were washed with 1x TBST for 5 minutes three times and then probed with Licor secondary antibody with 1:10,000 dilution in 1x TBST at room temperature for 1 hour.

To synchronize *Caulobacter* strains, cells were grown to an OD600 of 0.3-0.5 in PYE and swarmer cells were isolated using Percoll density centrifugation. Cells were released into PYE.

Flow cytometry was used to measure DNA content in rifampicin-treated cells as previously described (Chen et al., 2009). Cells were treated with rifampicin for three hours prior to fixing in 70% ethanol.

#### Carbon starvation

Cells were grown in M2G media (6.1 mM Na_2_HPO_4_, 3.9 mM KH_2_PO_4_, 9.3 mM NH_4_Cl, 0.5 mM MgSO_4_, 0.5 mM CaCl2, 10 μM FeSO_4_ (EDTA chelate), 0.2% glucose). Cells were then washed twice in cold M2G media (+glucose samples) or in M2 media (− glucose samples) and resuspended in pre-warmed M2G or M2 media.

#### Growth competition assay

Overnight cultures of CPC798 were mixed with either *NA1000* or *clpX*\* at a 1:1 ratio. For competition assays with *Δlon* strains, overnight cultures of CPC807 were mixed with either *Δlon* or *Δlon clpX*\* at a 1:1 ratio. Mixed strains were diluted into fresh PYE and allowed to outgrow for 12 doublings overnight. The initial populations were verified by phase contrast and fluorescent microscopy. Final ratios were normalized to their starting ratios. Data was plotted using GraphPad Prism.

#### Microscopy

Phase contrast images of logarithmically growing cells were taken by Zeiss AXIO Scope A1. Cells were mounted on 1% PYE agar pads and imaged using a 100X objective. Representative images of the same scale were cropped to display morphological defects.

#### Protein purification and modification

Untagged Lon, untagged ClpX and ClpX*, and his-tagged ClpP were purified as previously described (Chien et al., 2007b; Goldberg et al., 1994; Gur and Sauer, 2008b; Levchenko et al., 2000). Titin-I27-β20 was purified as previously described (Gur and Sauer, 2008b; Kenniston et al., 2003) and labeled with Fluorescein-5-maleimide (Thermo Scientific) under guanidine hydrochloride denaturation. The modified protein was buffer exchanged into H-buffer (20 mM Hepes pH 7.5, 100 mM KCl, 10 mM MgCl2, 10% glycerol) and stored at 4°C. The concentration of fluorescein-modified Titin-I27-β20 was determined using the Bradford assay (Thermo Scientific). FITC-Casein (Type III, Sigma) was prepared in water and stored at −20°C. His6SciP and his6CtrA were purified as described in (Gora et al., 2013b). DnaA and CcrM were purified as his6SUMO tagged proteins, followed by tag cleavage as described (Jonas et al., 2013; Liu et al., 2016; Wang et al., 2007). GFP-ssrA was purified as previously described (Yakhnin et al., 1998). CtrA-RD+15 was purified as described (Smith et al., 2014b). Detailed purification protocols are available upon request.

#### *In vitro* reconstitution assays

Degradation assays were performed at 30°C and monitored using SDS-PAGE gels as described previously (Bhat et al., 2013). The final concentrations used can be found in the figure legends.

#### Limited proteolysis

To perform limited chymotrypsin digestion, 0.004 mg/mL chymotrypsin was added to 0.01 mg/mL ClpX or ClpX* in the presence of ClpP and ATP regeneration system and incubated at 25°C in Hepes buffer (20 mM Hepes pH 7.5, 100 mM KCl, 10 mM MgCl2, 10% glycerol) for indicated time points. To quench the reactions, 5X SDS dye supplemented with 5 mM Protease inhibitor phenylmethylsulfonyl fluoride (PMSF) was added. The samples were run on a 10% Bis-Tris gel and transferred to nitrocellulose as described above. Membranes were probed with primary antibody in 3% milk in TBST at 4°C overnight with following dilution factors: 1:5000 dilution of ClpX. Membranes were washed with 1x TBST for 5 minutes three times and then probed with Licor secondary antibody with 1:10,000 dilution in 1x TBST at room temperature for 1 hour. The protein was visualized using Licor Odyssey CLx.

#### Binding of ADP to ClpX

625 nM mant-ADP was pre-incubated with 300 nM ClpX_6_/ClpX_6_* for 10 minutes. Subsequently, ATPγS was added to compete off the mant-ADP at various concentrations. Polarization measurements were read from 20 uL of these mixtures after an additional 15-minute incubation using opaque black NBS 384-well plates (Corning) and a SpectraMax M5 plate reader (Molecular Devices), with excitation and emission wavelengths set at 357 and 445 nm, respectively.

#### Differential scanning fluorimetry

Differential scanning fluorimetry was carried out in a Biorad realtime PCR thermocycler. 6 μM monomer of ClpX or ClpX* in the presence of 4 mM ATP or 12 μM ATP was mixed in a 1∶1 ratio with 4X Sypro Orange in Hepes Buffer (20 mM Hepes pH 7.5, 100 mM KCl, 10 mM MgCl_2_). The resulting mixtures were incubated at room temperature for 30 mins to allow the Sypro Orange to coat the proteins before subjecting triplicate samples to thermal unfolding from 25 °C to 95 °C. Melting temperatures were calculated by fitting of a 5-parameter sigmoid curve using the JTSA software (P. Bond, https://paulsbond.co.uk/jtsa).

#### Hydrogen-deuterium exchange mass spectrometry (HDX-MS)

Hydrogen-deuterium exchange experiments were carried out on a Synapt G2Si high-definition mass spectrometer (Waters) using an automated HDX robotics platform (Waters). Samples of 5.4 μM ClpX or ClpX* monomer were diluted 1:16 in D_2_O buffer containing 20 mM Hepes, 10 mM MgCl_2_, 100 mM KCl, and 5% glycerol. D_2_O and H_2_O buffers were supplemented with 4 mM ATP plus regeneration system (30 mM creatine kinase and 0.5 mg/mL creatine phosphate) for the high ATP condition and 12 μM ATP for the low ATP condition. Deuterium exchange was allowed to take place for 0, 10 seconds, 1 min, 10 minutes, and 60 min at 25 °C. The samples were subsequently quenched by addition of cold quench buffer (233 mM Sodium Phosphate pH 2.5) and run over an immobilized Waters ENZYMATE immobilized pepsin column (inner diameter: 2.1 × 30 mm) at a flow rate of 0.15 mL/min at high pressure (~11,000 psi) for peptide digestion. Blank runs were run in between each analysis to avoid peptide carry over. Continuous lock-mass correction was performed using leu-enkephalin compound. Peptides were ionized and separated by electrospray ionization for analysis at a mass resolution of 50 to 2,000 *m*/*z* range. Peptides were identified with triplicate undeuterated samples from each condition. Specific peptide regions were mapped onto the corresponding residues in the *E. coli* cryo-EM structure (PDB ID: 6PO1) using PYMOL (Version 2.5.0, Schrödinger). A model of ClpX in the potential open state was created by aligning each ClpX monomer to each ATPase monomer of the *Y. pestis* substrate-free cryo-EM structure (PDB ID: 6V11) by using the align function in PYMOL.

### QUANTIFICATION AND STATISTICAL ANALYSIS

Graphs were generated in GraphPad Prism. Unless otherwise stated in the figure legend, results represent the mean ± Standard Deviation (SD). Sample sizes are also reported in the figure notes. Error bars for the growth curves shown in Figure 1A represent the 95% confidence interval.

#### RNA sequencing data

Reads were mapped to the *Caulobacter* genome using BWA (Li and Durbin, 2009) and sorted with Samtools (Li et al., 2009).To obtain the number of reads per gene, *bedtools map* (Quinlan and Hall, 2010) was used. Rstudio and EdgeR was used to identify differentially expressed genes (R Core Team, 2019; Robinson et al., 2009).

Raw counts were normalized using the counts-per-million (CPM) method, and the *edgeR* package in R was used to perform differential gene expression analysis, KEGG pathway analysis, and FRY (Wu et al., 2010) gene set analysis. We filtered out any gene that had less than 30 normalized counts across 3 or more samples. To test for gene expression differences and identify differentially expressed genes across experimental groups, we used the quasi-likelihood F-test. The over-representation analysis for KEGG pathways done using the *kegga* function. FRY gene set analysis done using the *fry* function. The CcrM, DnaA, and SciP regulons were obtained from (Gonzalez et al., 2014; Hottes et al., 2005; Tan et al., 2010).

#### *In vivo* assays

For western blots, proteins were visualized using Licor Odyssey CLx. Bands were quantified using ImageJ (NIH) and degradation rates were plotted using Prism (Graphpad).

For bulk measurements of intracellular ATP, the Bactiter-Glo microbial cell viability assay kit (Promega) was used. Cells were mixed with an equal volume of BacTiter-Glo reagent in 384-well microtiter plates and incubated for five minutes before luminescence was measured.

For microscopy, MicrobeJ (Ducret et al., 2016) for ImageJ (Schneider et al., 2012) was utilized to quantify cell lengths. Stalk lengths were quantified using ImageJ. Prism was utilized for representations of cell and stalk length measurements.

#### *In vitro* assays

Densitometry for degradation assays was performed with ImageJ and degradation rates were plotted using Prism.

Degradation of FITC-Casein and FT-Titin-B20 was monitored as an increase in fluorescence over time. GFP-CtrARD+15 degradation was observed as a loss of fluorescence over time as described previously (Smith et al., 2014b).

ATP hydrolysis for ClpXP and ClpX*P was measured using a coupled kinase assay as previously described (Burton et al., 2003).

For binding of ADP to ClpX, IC50s were calculated by fitting the polarization data using GraphPad Prism to an inhibitor versus response – variable slope (four paramaters) function in GraphPad Prism: Y=Bottom + (Top-Bottom)/(1+(IC50/X)^HillSlope) where IC50 is the ATPγS concentration halfway between bottom and top.

For differential scanning fluorimetry, melting temperatures were calculated by fitting of a 5-parameter sigmoid curve using the JTSA software (P. Bond, https://paulsbond.co.uk/jtsa).

#### HDX-MS data

HDX-MS experiments were performed in triplicate. Identification of peptides and analysis of the uptake plots and charge states for each peptide were completed in Protein Lynx Global Server and DynamX (v. 3.0, Waters). The following processing paramaters were utilized: minimum peptide intensity of 1000, minimum peptide length of 5, maximum peptide length of 30, minimum MS/MS products of 2, minimum products per amino acid of 0.25, minimum score of 5, and maximum MH+ error threshold of 15 p.p.m. Woods plots were generated by Deuteros (Lau et al., 2021) using the peptide significance test (p-value <0.01). HDX data summary for each condition was exported from Deuteros and can be found in Supplementary File 3. No correction was made for back-exchange.

#### Descriptive titles for Supplemental Files

DataS1. This file contains the RNA sequencing experiments referred to in Main Figure 1. DataS1.xlsx

DataS2. This file contains the Hydrogen Deuterium Exchange log for experiments shown in Main Figure 6. DataS2.pdf

## Notes

https://www.ncbi.nlm.nih.gov/geo/query/acc.cgi?acc=GSE166765

### Competing Interest Statement

The authors have declared no competing interest.

### Summary of Updates

This version of the manuscript includes additional experiments such as Hydrogen-deuterium exchange mass spectrometry and differential scanning fluorometry that were performed to shed light on the mechanisms that govern protease substrate specificity.

