## Supplementary File 2: HDX Log File for "ATP hydrolysis tunes specificity of a AAA+ protease"

Supplemental File 2- HDX Log file

State A: ClpX_CAUVN (clpx sat atp)
HDX time course (min): 0, 0.17, 1, 10, 60
HDX control samples: None
Back-exchange (mean / IQR): N/A
### of Peptides: 129
Sequence coverage: 93.10%
Average peptide length / redundancy: 13.07 / 4.31
Replicates (technical): 3
Repeatability: 0.1915 (average SD)

State B: CLPX_CAUVN (clpxg178a sat atp)
HDX time course (min): 0, 0.17, 1, 10, 60
HDX control samples: None
Back-exchange (mean / IQR): N/A
### of Peptides: 125
Sequence coverage: 92.62%
Average peptide length / redundancy: 13.35 / 4.29
Replicates (technical): 3
Repeatability: 0.2475 (average SD)

State B: CLPX_CAUVN (clpx 12um atp)
HDX time course (min): 0, 0.17, 1, 10, 60
HDX control samples: None
Back-exchange (mean / IQR): N/A
### of Peptides: 127
Sequence coverage: 93.10%
Average peptide length / redundancy: 13.23 / 4.30
Replicates (technical): 3
Repeatability: 0.3069 (average SD)
