## Supplementary Figures for "ATP hydrolysis tunes specificity of a AAA+ protease"

#### Supplementary Figure Legends

##### Figure S1: Validation of motility screen suppressor

(A-B) Motility assay on 0.3% PYE agar of indicated strains.
C. Sequence alignment of ClpX homologs. Conserved glycine residue and walker B motif are marked.
D. Quantification of the percent of cells with DNA content of 1N, 2N, and >2N in *wt*, *Δlon,* and *Δlon clpX** strains.
E. Competition assay with Δ*lon* cells harboring xylX::Plac-venus (constitutive venus expression) and nonfluorescent Δ*lon* or Δ*lon clpX** strains. Exponential phase cells were mixed 1:1, diluted, and allowed to outgrow for 12 doublings. Quantification of triplicate experiments is shown below. Error bars represent SD.

##### Figure S2: RNA-seq gene set analysis

(A-B) Volcano plot showing the transcriptional differences of Δ*lon* compared to *wt* (left) and *Δlon clpX** compared to *wt* (right), as measured by RNA-seq. The negative $\log_{10}$ of the p value is plotted against $\log_2$ of the fold change mRNA counts. Genes that are marked in red have an FDR <0.05. Transcripts that are associated with either the SciP regulon (Tan et al., 2010) or cell cycle (KEGG pathways) are highlighted in blue.
C. Kyoto Encyclopedia of Genes and Genomes (KEGG) pathway enrichment analysis and FRY gene set analysis. FRY gene set analysis performed using *fry* function from *edgeR* package in R (Robinson et al., 2009; Wu et al., 2010). The CcrM, DnaA, and SciP regulons were assigned according to (Gonzalez et al., 2014; Hottes et al., 2005; Tan et al., 2010). Over-represented pathways in KEGG pathway database identified using the *kegga* function from *edgeR* package in R.
D. Western blot showing CcrM levels in synchronized populations of wt, *Δlon, and Δlon clpX** cells. Swarmer cells were isolated using a density gradient and an equal number of cells were released into fresh PYE medium. Samples were withdrawn at the indicated time points and probed with anti-CcrM.
E**.** Growth curves of *wild type* (wt), *Δlon, and Δlon* merodiploid with second copy of *clpX** at native locus*.* Cells grown in PYE.
F. Spot assays comparing colony formation of strains in PYE and PYE supplemented with mitomycin C.
G. Motility assay on 0.3% PYE agar of indicated strains

##### Figure S3: *In vitro* characterization of ClpX*

A. ATPase activity of 0.1 μM $ClpX_6$/$ClpX^*_6$ plus 0.2 μM $ClpP_{14}$ is displayed. Triplicate experiments are shown.
B. *In vitro* degradation of CcrM by 0.2 μM $Lon_6$ is shown as a control. The bottom gel shows CcrM degradation assay in the presence of ATP, with no regeneration mix.
C. FITC-Casein titration in the presence of either ClpXP or ClpX*P. Data was fit to the Michaelis-Menten equation. ClpX*P degrades FITC-Casein 2-3 fold better than ClpXP.

F. Fluorescin-*Titin*-I27-β20 titration in the presence of either ClpXP or ClpX*P. Data was fit to the Michaelis-Menten equation. $V_{max}/K_M$ was 30% higher for ClpX*P in comparison to ClpXP. Inset shows residual plot for Michaelis-Menten fit.

**Figure S4: ClpX*P is deficient in native substrate degradation**
A. *In vitro* degradation of GFP-ssrA.
B. *In vitro* fluorescence degradation assay of eGFP-CtrA-RD+15 in the presence of adaptors. Degradation assays were performed with 1 μM eGFP-CtrA-RD+15, 0.1 μM $ClpX_6$ or $ClpX^*_6$, 0.2 μM $ClpP_{14}$, 2 μM CpdR, 1 μM RcdA, 1 μM PopA, and 20 μM cyclic di-GMP, and ATP regeneration system.

**Figure S5: Modulating ATP changes substrate preferences**
A. Titration of ATP against constant mant-ADP (1 μM) in the presence of either 300 nM $ClpX_6$ or $ClpX^*_6$ assayed by changes in fluorescence. Data was fit to an inhibitor versus response – variable slope (four parameters) function in Graphad Prism:
Y=Bottom + (Top-Bottom)/(1+(IC50/X)^HillSlope) where IC50 is the ATP concentration halfway between bottom and top. Two independent replicates shown.
B. ATPγS competition assay in the presence of either 300 nM $ClpX_6$ or $ClpX^*_6$. Fluorescence polarization of 625 nM mant-ADP was measured as described in methods. Data was fit as described in A. Two independent replicates shown.
C. GFP-ssrA degradation by ClpXP or ClpX*P as a function of ATP concentration. Assays were performed with 0.1 μM $ClpX_6$ or $ClpX^*_6$, 0.2 μM $ClpP_{14}$, ATP regeneration system, and 10 μM GFP-ssrA.
D. *In vitro* degradation of SciP by ClpXP under low and saturating ATP conditions. Assays were performed with 0.1 μM $ClpX_6$, and 0.2 μM $ClpP_{14}$ and 5 μM SciP. Quantification of triplicate experiments shown.
E. Relative intracellular ATP concentrations measured using a luciferase-based assay during the course of antibiotic shutoff assay.

**Figure S6: ClpX* mutation and limiting ATP lead to increased dynamics of ClpX**
A. Woods plot comparing deuterium uptake for wildtype ClpX at 4 mM ATP vs 12 μM ATP at 60 mins. Each bar on Woods plot represents a single peptide with peptide length corresponding to the bar length. Red bars indicate a deprotected (more deuterium uptake) region, blue represents a protected region, and gray bars are not significantly different. Woods plots were created with Deuteros (Lau et al. 2021) using the peptide significance test (p-value <0.01).
B. Woods plot comparing deuterium uptake for wildtype ClpX vs ClpX* at 4 mM ATP at 0.17, 1, and 10 minutes. Each bar on Woods plot represents a single peptide with peptide length corresponding to the bar length. Red bars indicate a deprotected (more deuterium uptake) region, blue represents a protected region, and gray bars are not significantly different. Woods plots were created with Deuteros using the peptide significance test (p-value <0.01).
C. Comparison of deuterium uptake plots of selected peptides for wildtype ClpX vs ClpX*.

D. Each ClpX monomer (Protein Data Bank ID: 6PO1) was mapped onto the substrate-free Lon ATPase domain (Protein Data Bank: 6V11) using PYMOL. Residues 186-200 highlighted in red.
E. We propose that ClpX exists in an equilibrium between a 'closed' and 'open' set of conformations and promoting one state over the other leads to alterations in substrate specificity. In the presence of ClpX* or in ATP-limited conditions, ClpX adopts a more open conformation, allowing capture and recognition of substrates such as casein. The balance shifts to the closed state under high ATP conditions, allowing degradation of substrates such as GFP-ssrA, which preferentially bind the closed state.

Figure S1 - related to figure 1

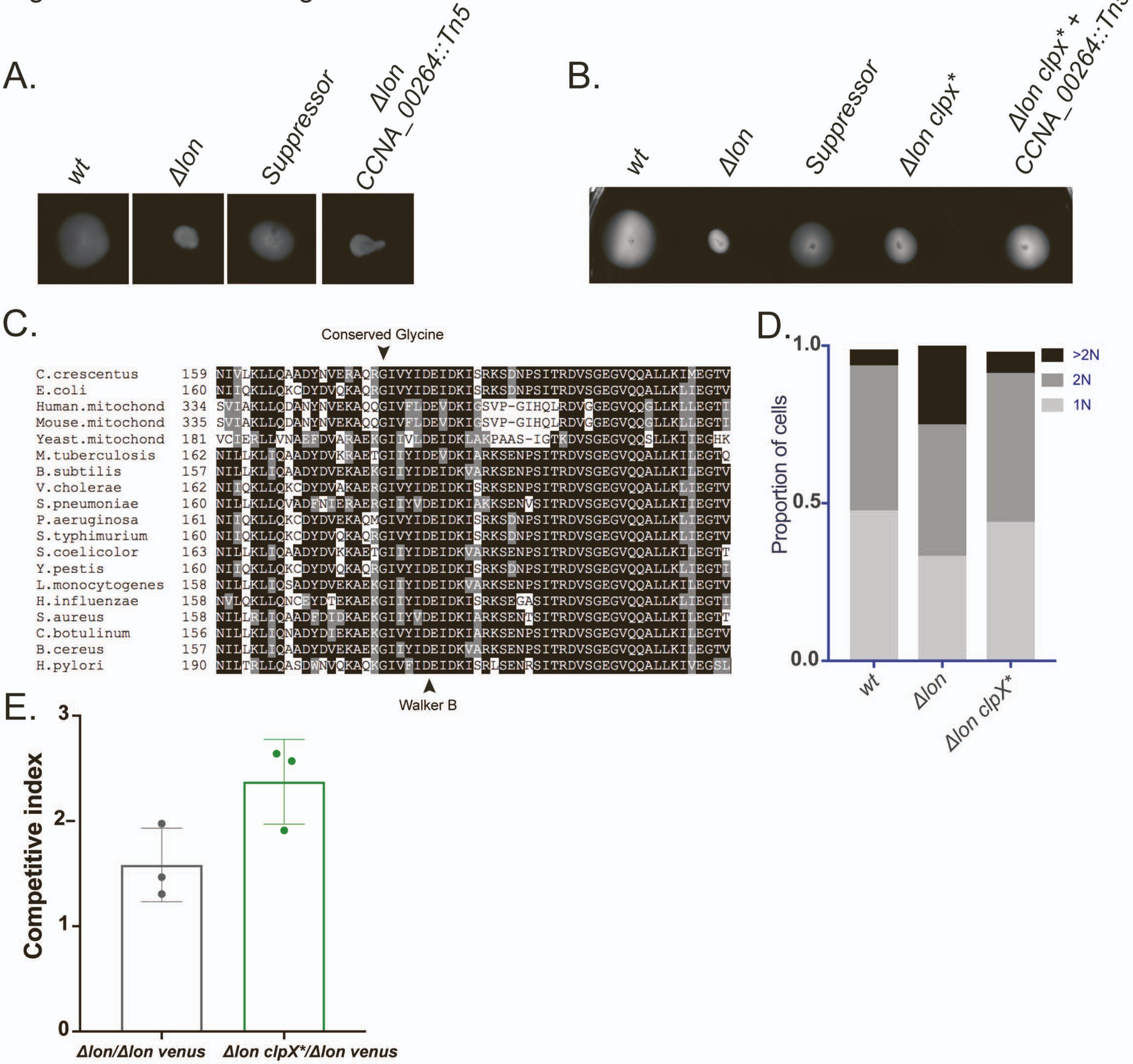

Figure S2- Related to Figure 2

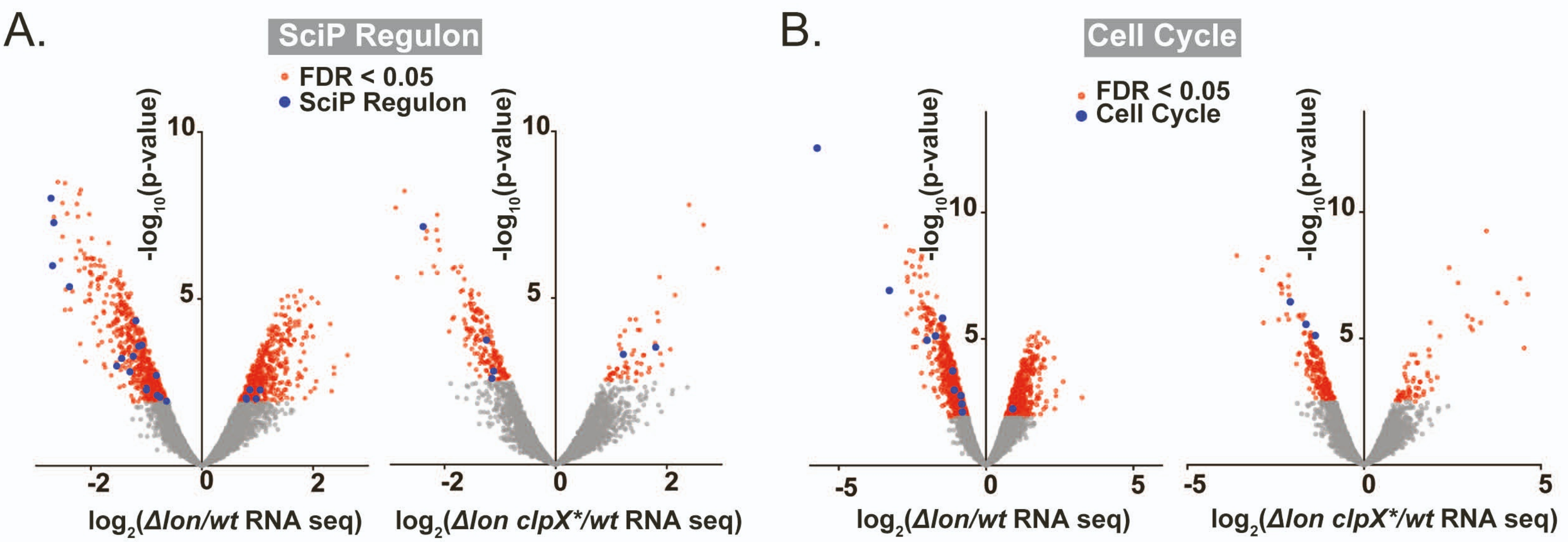


C.

##### FRY Gene Set Analysis

| | *Δlon* vs *wt* | | | | Δlon clpX* vs wt | | | |
|---|---|---|---|---|---|---|---|---|
| | Number of Genes | Direction | p-value | FDR | Number of Genes | Direction | p-value | FDR |
| SciP | 61 | Down | 0.0029 | 0.0044 | 61 | Down | 0.3391 | 0.5042 |
| CcrM | 504 | Up | 0.0014 | 0.0014 | 504 | Up | 0.7391 | 0.7391 |
| DnaA | 32 | Down | 0.0989 | 0.0980 | 32 | Down | 0.0919 | 0.1275 |

##### KEGG Enrichment Analysis

| | Cell Cycle | | |
|---|---|---|---|
| | Number of Genes | Direction | p-value |
| *Δlon vs wt* | 31 | 10, Down | 0.0086 |
| *Δlon clpX* vs wt* | 31 | 3, Down | 0.1725 |

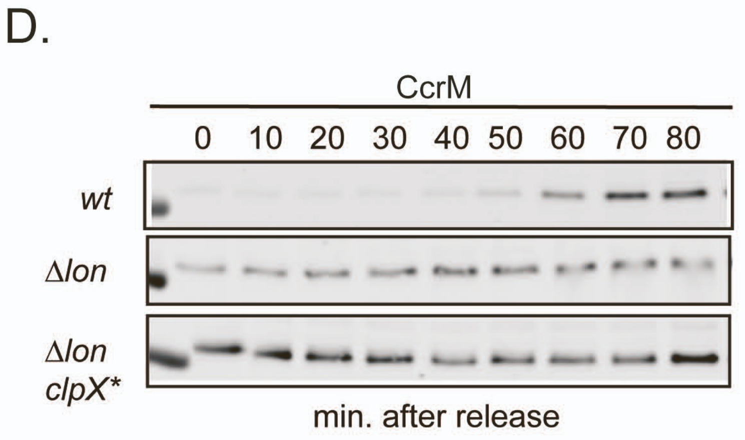


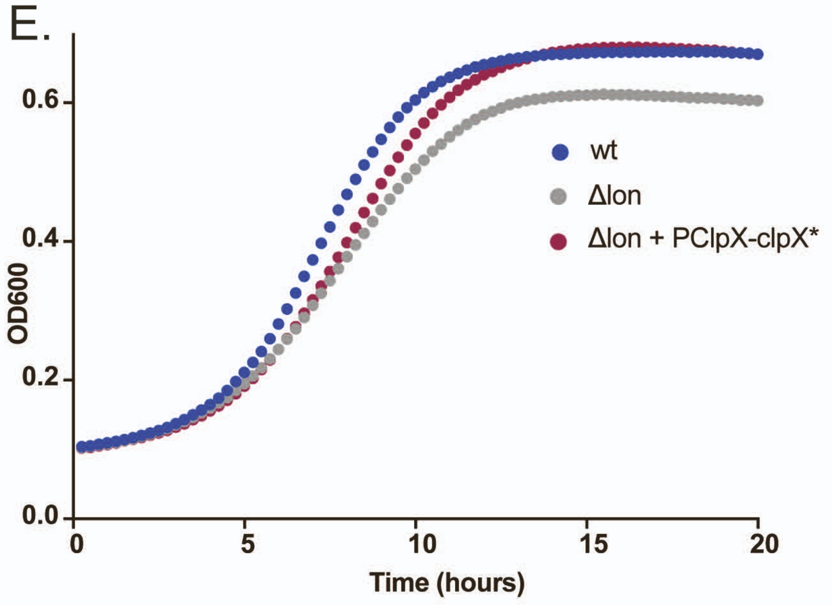


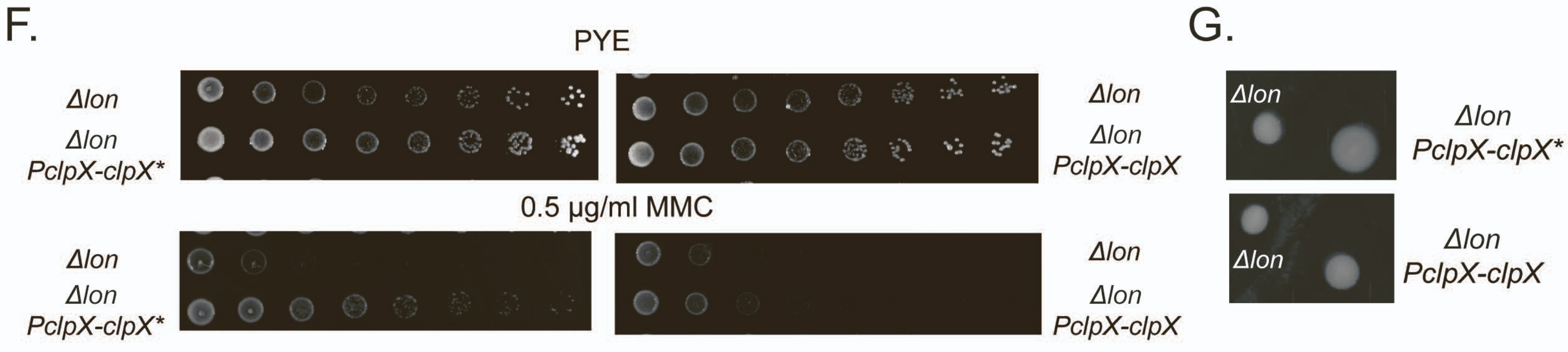

### Figure S3- Related to Figure 3

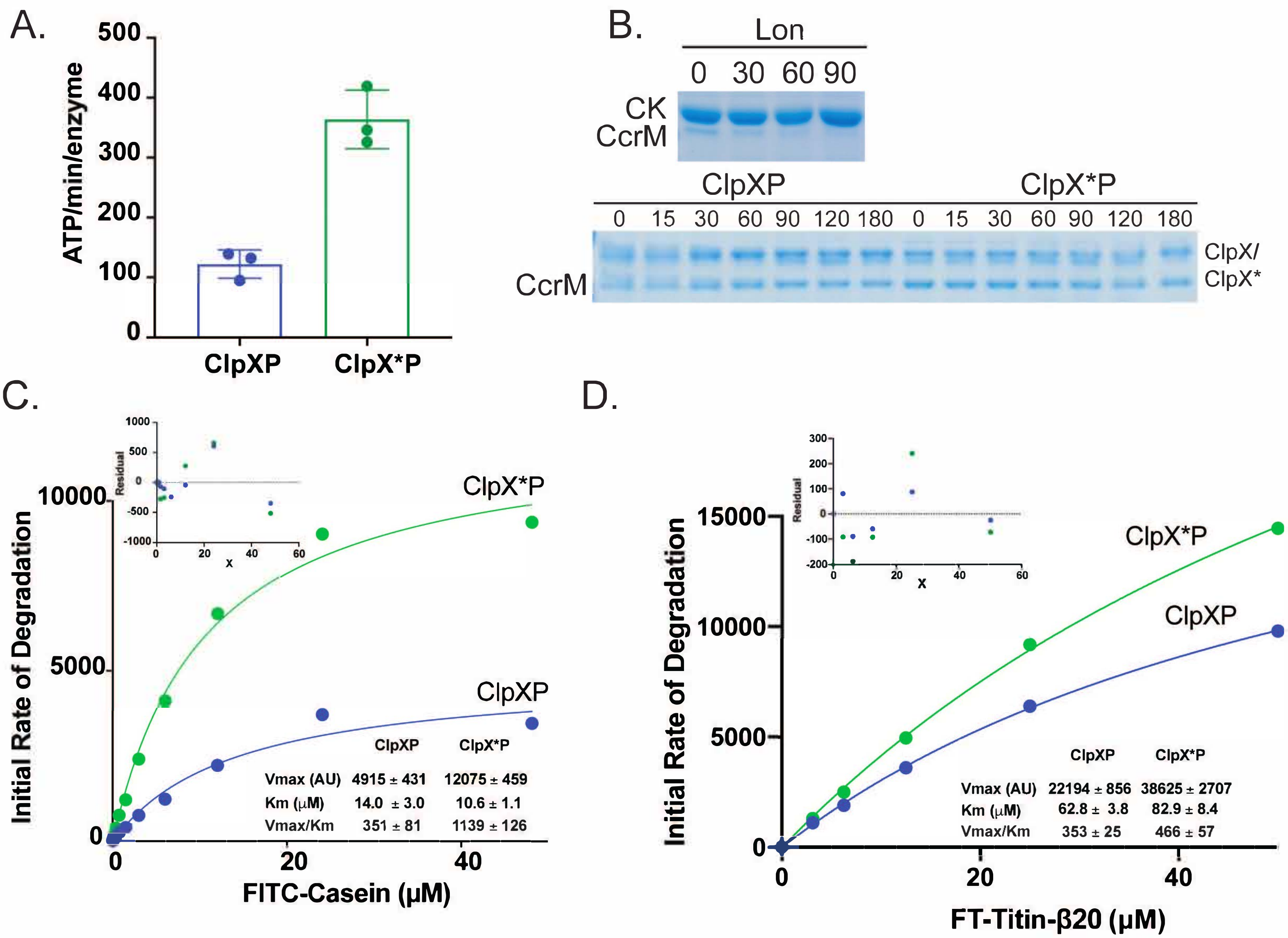

Figure S4- Related to Figure 4

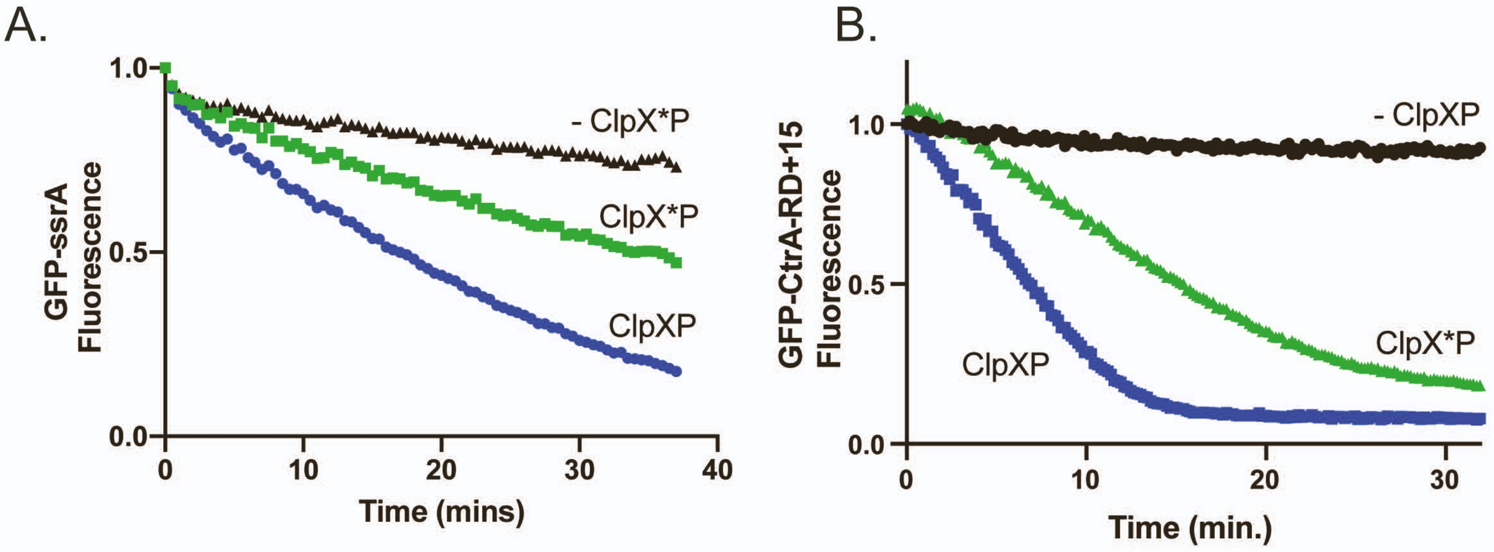

Figure S5- Related to Figure 5

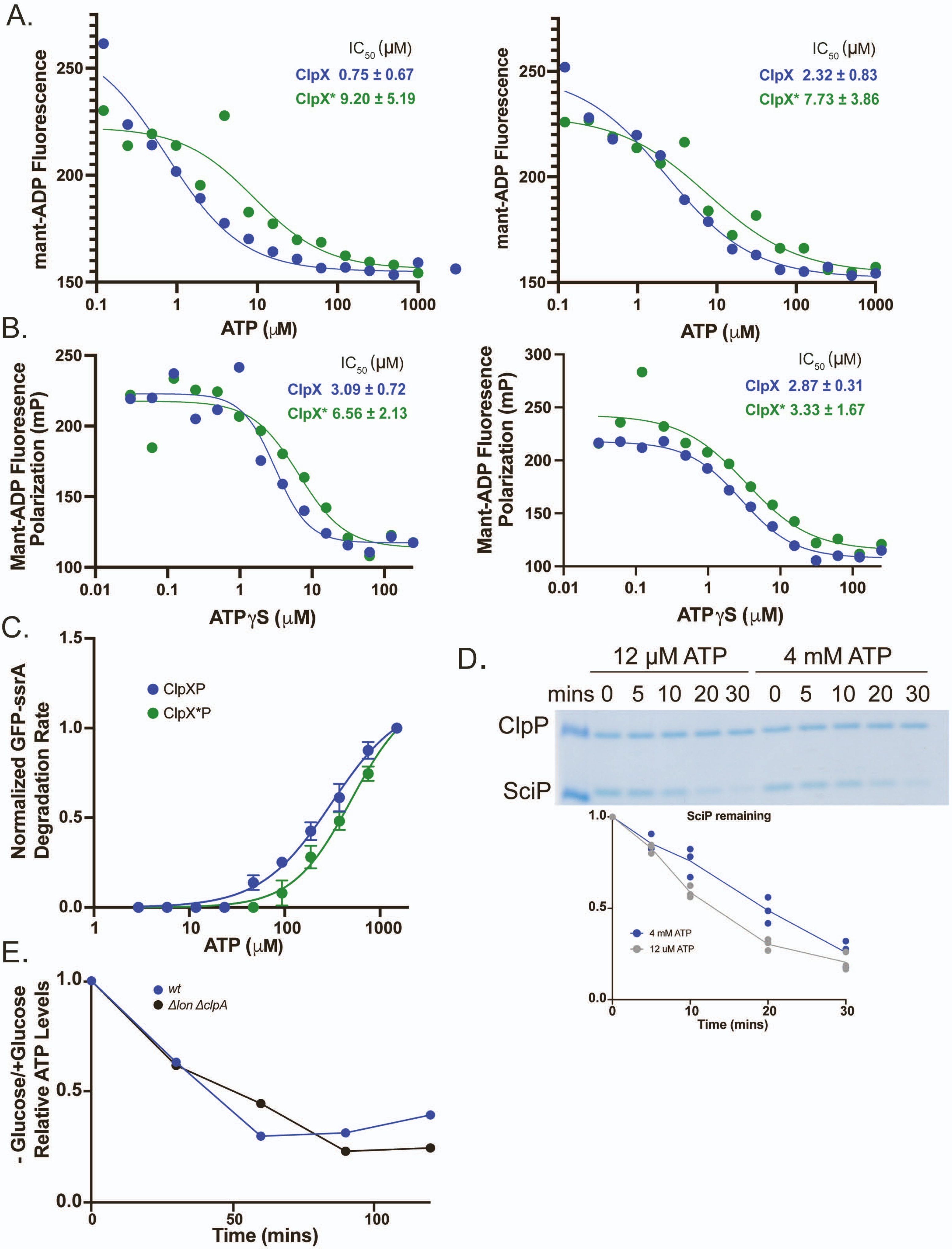

### Figure S6- Related to Figure 6

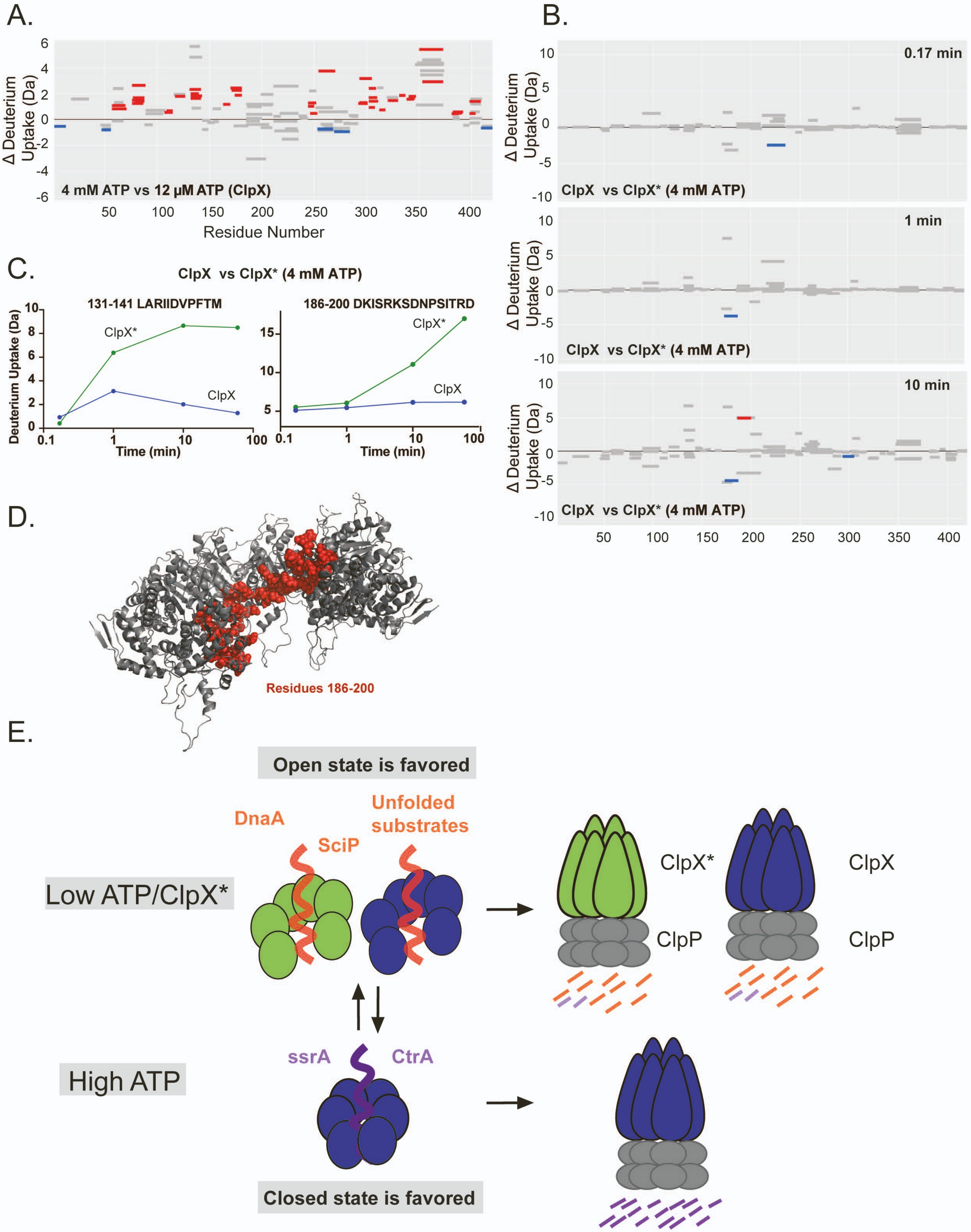
